# Epigenetic silencing by SMYD3 represses tumor intrinsic interferon response in HPV-negative squamous cell carcinoma of the head and neck

**DOI:** 10.1101/2022.11.04.515098

**Authors:** N Nigam, B Bernard, S Kim, K Burkitt, D Tsai, Y Robbins, C Sievers, A Clint, RL Bennett, TT Tettey, B Carter, R Bao, L Rinaldi, M.W. Lingen, H Sater, EF Edmondson, H Cheng, X Luo, K Brennan, C Chen, S Sevilla, M Murali, S Sakata, K Takeuchi, Y Nakamura, R Uppaluri, JB Sunwoo, C Van Waes, JD Licht, GL Hager, V Saloura

**Author notes:** Corresponding author and lead contact: Vassiliki Saloura, MD, PhD, 41 Medlars Drive, Bethesda, MD, 20892, Thoracic and GI Malignancies Branch, National Cancer Institute. These authors contributed equally (N Nigam, B Bernard).

## Abstract

Cancers often display immune escape, but the mechanisms and potential for reversibility are incompletely understood. Epigenetic dysregulation has been implicated in the immune escape of various cancer types. We have identified the epigenetic modifier SET and MYND-domain containing protein 3 (SMYD3) as a mediator of immune escape in human papilloma virus (HPV)- negative head and neck squamous cell carcinoma (HNSCC), an aggressive disease with poor prognosis and low response to immunotherapy with pembrolizumab, a programmed-death-1 (PD-1) targeting antibody. SMYD3 loss increased the sensitivity of HNSCC cancer cells to IFN-β, resulting in upregulation of type I IFN response and antigen presentation machinery genes. We found that SMYD3 regulates the transcription of Ubiquitin-Like PHD And RING Finger Domain-Containing Protein 1 (UHRF1), a key epigenetic reader of trimethylated lysine 9 on histone H3 (H3K9me3), which binds to H3K9me3-enriched promoters of key immune-related genes and silences their expression. SMYD3 further maintains the repression of immune-related genes through the deposition of H4K20me3 within the gene body regions of these genes. In an anti-PD-1 immune checkpoint resistant syngeneic mouse model of HPV-negative HNSCC, Smyd3 depletion induced influx of CD8+ T-cells, upregulated PD-L1 and MHC class I molecules, and increased sensitivity to anti-PD-1 therapy. SMYD3 overexpression was associated with decreased CD8 T-cell infiltration in tumor samples from patients with HPV-negative HNSCC, and was associated with poor response to pembrolizumab. Overall, these data highlight a previously unreported function of SMYD3 as a master epigenetic regulator of anti-tumor immune response in HPV-negative HNSCC and provide a rationale for translational approaches combining SMYD3 depletion strategies with checkpoint blockade to overcome anti-PD-1 resistance in this devastating disease.

## INTRODUCTION

Head and neck squamous cell carcinoma (HNSCC) affects approximately 50,000 patients annually in the United States (Siegel et al., 2020). While human papilloma virus (HPV)-positive patients tend to have an excellent prognosis (Ang and Sturgis, 2012), HPV-negative patients have an approximately 50% recurrence rate after treatment with standard cisplatin-based chemoradiotherapy and/or surgery. Furthermore, patients with recurrent/metastatic (R/M) HNSCC treated with platinum-based chemotherapy regimens have a dismal prognosis with a median overall survival of approximately 10 months (Chaturvedi et al., 2011).

Recently, pembrolizumab, which blocks the programmed-death-1 (PD-1)/programmed death-ligand 1 (PD-L1) axis, was approved as a first-line therapy for patients with R/M HNSCC (Burtness et al, 2019). However, response rates in HPV-negative HNSCC patients are as low as 19%, underscoring the presence of immune escape mechanisms. While multiple clinical trials are currently attempting to reduce or overcome immune suppressive mechanisms in the tumor microenvironment (Ferris et al, 2015), the molecular factors involved in CD8+ T-cell exclusion in HPV-negative HNSCC are still not well understood. Particularly in HPV-negative disease, ∼80% of tumors demonstrate low CD8+ T-cell infiltration (Saloura et al, 2019), which is an independent predictor of poor response to PD-1 inhibition (Hanna et al, 2018). Thus, the elucidation of mechanisms that dictate poor CD8+ T-cell infiltration in HPV-negative HNSCC is of paramount importance to improve immunotherapy-based treatment strategies (Bonaventura et al 2019, Duan et al 2020).

Epigenetic regulation mediated by histone modifications has emerged as an important therapeutic avenue for anticancer therapy. The Cancer Genome Atlas (TCGA) has documented a plethora of genetic and expression alterations in chromatin modifiers in multiple cancer types, including HPV-negative HNSCC (Gao et al, 2013, Cerami et al, 2012, Leemans et al 2018). A significant body of preclinical evidence supports their oncogenic function not only as epigenetic regulators through histone modification, but also through non-histone protein modifications (Hamamoto et al, 2015). In addition, recent evidence supports that some of these chromatin modifiers may be involved in the regulation of antitumor immunity (Cao et al, 2020).

Given the importance of immunotherapy in HPV-negative HNSCC, we recently conducted a systematic bioinformatic interrogation of two large HPV-negative HNSCC expression cohorts to identify protein methyltransferases (PMTs) and demethylases (PDMTs), a class of chromatin modifiers that are frequently genetically altered in HPV-negative HNSCC, that may be associated with the non-inflamed phenotype and could thus be biological culprits of CD8+ T-cell exclusion in this disease (Vougiouklakis et al, 2017). We found that the mRNA expression levels of Set and MYND domain protein 3 (SMYD3), a protein lysine methyltransferase (PMT) with oncogenic effects in several cancer types, correlated inversely with the mRNA levels of CD8A, CD8+ T-cell attracting chemokines, such as CXCL9, CXCL10 and CXCL11, as well as a panel of antigen presentation machinery (APM) molecules, such as TAP1, TAP2, TAPBP, and HLA-A/B/C and B2M. Additionally, SMYD3 knockdown in two HPV-negative HNSCC cell lines in vitro led to significant upregulation in mRNA and protein levels of CXCL9, CXCL10 and CXCL11, suggesting a role of SMYD3 in regulating the transcription of these genes.

SMYD3, a member of the SET and MYND-domain family, is a PMT that has been implicated as an oncogene in multiple cancer types, such as colorectal cancer, hepatocellular carcinoma (Sarris et al, 2016, Bernard et al, 2021), K-ras mutant pancreatic ductal and lung adenocarcinoma (Mazur et al, 2014), and breast cancer (Hamamoto et al 2004, Wang et al, 2019, Fenizia et al 2019). In hepatocellular carcinoma, SMYD3 acts predominantly as a chromatin modifier in the nucleus and exerts its oncogenic activity by trimethylating and reading histone 3 lysine 4 (H3K4me3) (Sarris et al, 2016). SMYD3 binds directly to DNA as a transcription factor, forms a complex with RNA polymerase II and reads H3K4me3 to transactivate a set of oncogenes involved in cell cycle and epithelial-mesenchymal transition (Sarris et al, 2016). A less studied histone substrate of SMYD3 is H4K20, which has been reported to be trimethylated by SMYD3 and induces transcriptional repression (Foreman et al, 2011, Jiang et al, 2019). However, in K-ras mutant pancreatic and lung adenocarcinomas, SMYD3 directly methylates the non-histone, cytoplasmic substrate MAP3K2, inhibiting its interaction with phosphatase PP2A and increasing the K-ras to Erk1/2 downstream signaling pathway (Mazur et al, 2014). This suggests that the oncogenic function of SMYD3 may vary by cancer type and may be mediated either through its function as a chromatin modifier/transcription factor or as a direct methylator of cytoplasmic substrates.

SMYD3 inhibitors are actively in development (Peserico et al, 2015, Mitchell et al, 2015, Huang et al, 2019), augmenting siRNA and CRISPR as tools for investigation of the mechanism and therapeutic potential of SMYD3. Recently, Kontaki et al (Kontaki et al, 2021) showed that next generation anti-sense oligonucleotides (ASOs) targeting Smyd3 decreased Smyd3 mRNA levels *in vivo* efficiently and halted the growth of liver tumors in a chemically induced HCC mouse model. ASOs are a promising drug discovery platform of RNA-targeted therapeutics involving single-stranded, chemically modified DNA oligonucleotides, and constitute an alternative drug platform over enzymatic inhibitors or antibodies (Crooke et al, 2018).

In the present study, we demonstrate that genetic depletion of SMYD3 by siRNAs, SMYD3 ASOs or CRISPR transcriptionally derepresses the expression and leads to an orchestrated induction of multiple type I IFN response and APM genes in response to IFN-β in HPV-negative HNSCC cells. Transcriptional repression of these genes is dependent on SMYD3-mediated transcriptional upregulation of Ubiquitin-Like PHD And RING Finger Domain-Containing Protein 1 (UHRF1), a key epigenetic regulator that reads the repressive trimethylated lysine 9 on histone H3 (H3K9me3) mark, and on the deposition of H4K20me3 on immune-related genes. Systemic Smyd3 ASO treatment in an anti-PD-1 resistant immunocompetent mouse model of flank MOC1 tumors induces influx of CD8+ T-cells, upregulates PD-L1 and MHC class I molecules, and increases sensitivity to anti-PD-1 therapy. Finally, SMYD3 is overexpressed in HPV-negative HNSCC tumors and baseline SMYD3 and UHRF1 mRNA tumor expression levels predict response to anti-PD-1 neoadjuvant therapy in HPV-negative HNSCC patients. These data identify SMYD3 as a master epigenetic regulator of anti-tumor immune response in HPV-negative HNSCC, and provide a rationale for translational approaches combining SMYD3 ASOs with checkpoint blockade to overcome anti-PD-1 resistance in this devastating disease.

## RESULTS

### SMYD3 depletion is associated with upregulation of type I IFN response and APM genes in HPV-negative HNSCC cells

Based on our previously published work (Vougiouklakis T et al, 2017), we sought to evaluate whether SMYD3 depletion has a systemic effect on the expression of type I IFN response and APM genes (here on referred to cumulatively as immune-related genes, see Statistical Analyses, “Lists of type I IFN response and APM genes”) (**Supplementary Tables 1, 2**).

To this purpose, we conducted RNA-sequencing (seq) analysis of the HPV-negative HNSCC cell line HN-6 with endogenously high levels of SMYD3 before and after siRNA-mediated SMYD3 knockdown for 72h and after 24h of IFN-β exposure. Evaluation of the expression levels of immune-related genes revealed upregulation of multiple key type I IFN response genes, such as *MX1*, *MX2*, *CD274*, *IL1B*, *GBP2*, *IL6* (*IFN-β*), and APM genes, such as *TAP2*, *TAPBP*, *B2M*, and *HLA-A* and *-B* (**Fig.1A**, **Supplementary Fig.1A, B**). This phenotype was reproduced with pharmacological targeting of SMYD3 using SMYD3 ASOs (Kontaki et al, 2021, Ionis Pharmaceuticals) (**Fig.1B, Supplementary Fig.2A, B)**. Consistently, the RNA-seq profile of a SMYD3 CRISPR knockout (KO) cell line (termed 5-3) was compared with those of parental HN-6 cells, and exhibited a similar phenotype (**Fig.1C, Supplementary Fig.3 A, B)**. More specifically, SMYD3 depletion in the SMYD3 KO cell line induced significant upregulation of 73 of 97 IFNa response genes and 24 of 88 APM genes (**Fig. 1D, Supplementary Table 2**). Among the three cell systems, 28 of 97 IFNa response genes were commonly and significantly upregulated (**Supplementary Fig. 4, Supplementary Fig. 5**). Concordantly, Gene Set Enrichment Analysis (GSEA) revealed enrichment in pathways related to inflammation in all three cell systems (**Fig.1E, Supplementary Fig. 6**).

**Figure 1.**
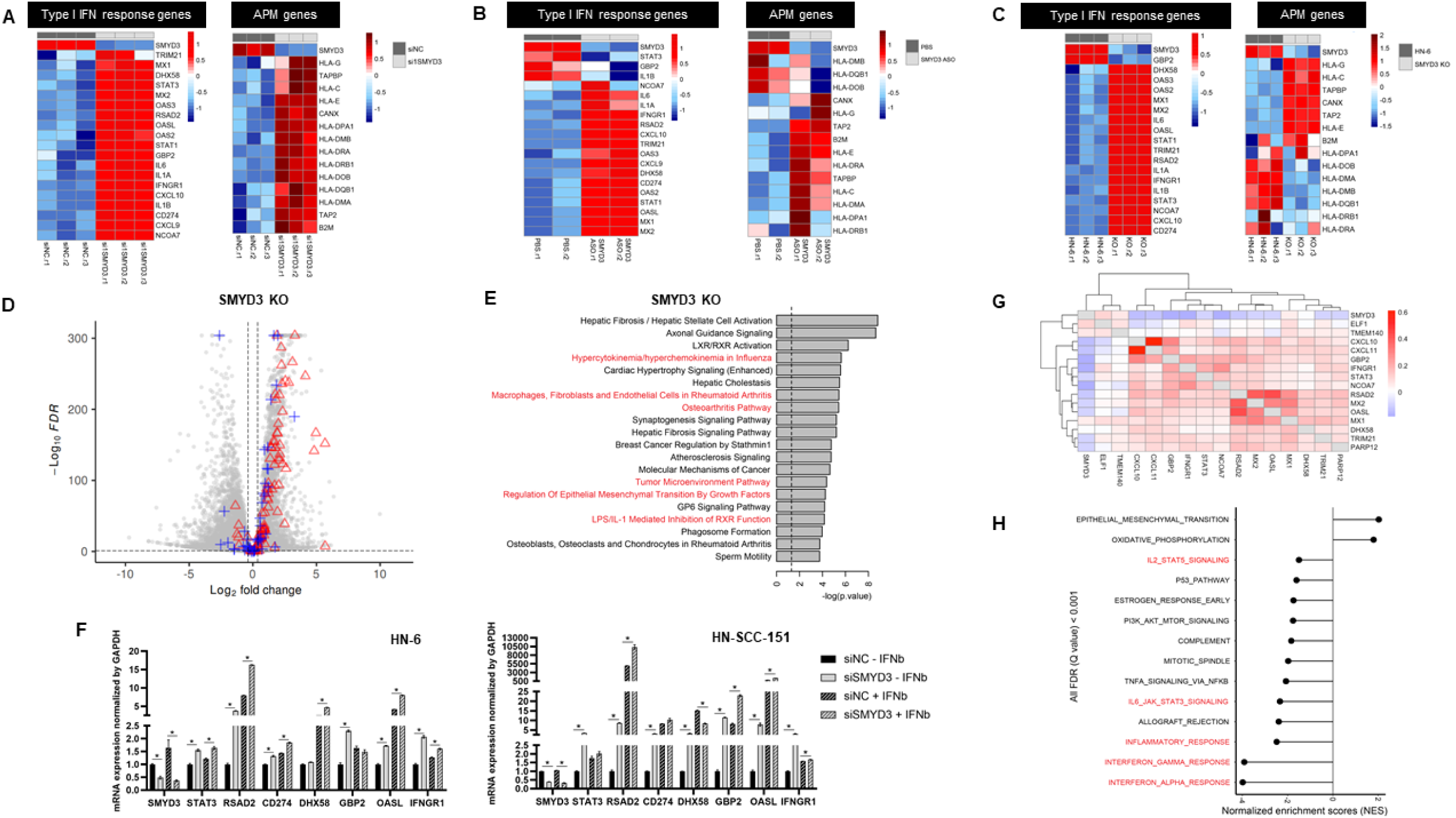
SMYD3 depletion induces upregulation of type I IFN response and antigen presentation machinery (APM) genes in HPV-negative HNSCC cells. **(A, B, C).** Heatmaps of type I IFN response and APM genes (RNA-seq) in HN-6 cells after SMYD3 depletion. Heatmaps using z-score of variance stabilizing transformed expression values. **(A)** HN-6 cells treated with control siRNAs or a SMYD3-targeting siRNA for 72h (three biological replicates per condition, siNC.r : control replicate, siSMYD3.r: SMYD3 siRNA replicate), and exposed to 1000U/ml of IFN-β for 24h (day 2 of transfection), **(B)** HN-6 cells treated with PBS or SMYD3 ASOs for 72h (three biological replicates per condition), and exposed to 1000U/ml of IFN-β for 24h (day 2 of ASO treatment), **(C)** parental HN-6 and SMYD3 knockout (KO) cells (5-3 cell line) after exposure to 1000U/ml of IFN-β for 24h. **(D)** Volcano plot showing DESeq2 results in SMYD3 KO cells (5-3 cell line) compared to parental HN-6 cells. FDR < 0.1, log2FC threshold: log2 (1.3). Red triangles: IFNa genes (from GSEA gene set, upregulated: 73, downregulated: 7), blue crosses: APM genes (from GSEA gene set, upregulated: 24, downregulated: 11), gray circles: other genes. Total number of genes=19,447 (upregulated: 6,300, downregulated: 7,292). **(E)** Gene Set Enrichment Analysis (GSEA) reveals enrichment of pathways related to inflammation in an HPV-negative cell line (HN-6) after SMYD3 knockout (KO, 5-3 cell line). **(F)** Quantitative PCR (SYBR green) for immune-related genes in HN-6 (left panel) and HN-SCC-151 cells (right panel) after siRNA-mediated SMYD3 knockdown. Cells were treated with control siRNAs and a SMYD3-targeting siRNA for 72h with or without exposure to IFN-β (1000U/ml) for 24h (day 2 of transfection). **(G)** Pairwise correlations between SMYD3 and type I IFN response genes in single cancer cells of a publicly available single-cell RNA-seq database. **(H)** Normalized enrichment scores of Hallmark gene sets correlated with SMYD3 expression. Positive NES indicates activation of the pathway in the presence of SMYD3. Negative NES indicates suppression of the pathway in the presence of SMYD3. The Cancer Genome Atlas, Firehose Legacy, 427 HPV-negative tumors samples.

To confirm these findings and further evaluate whether the effect of SMYD3 on the expression of immune-related genes is dependent on IFN-β exposure, we treated two HPV-negative HNSCC cell lines (HN-6, HN-SCC-151) with negative control or SMYD3-specific siRNAs for 72h with or without exposure to IFN-β, and conducted qPCR for a panel of representative type I IFN response genes. Results confirmed that SMYD3 depletion induced significant upregulation of multiple immune-related genes in both cell lines (**Fig.1F**). Importantly, significant upregulation of these genes was observed even in the absence of IFN-β. This suggests that the effect of SMYD3 on the expression of some immune-related genes is not IFN-β dependent, and thus, SMYD3 depletion could trigger an inflammatory response in cancer cells even in the setting of a “cold” tumor microenvironment.

To evaluate the correlation between SMYD3 and the expression of type I IFN response and APM genes in single cancer cells from patient tumor samples, we analyzed a publicly available single-cell RNA-seq database of HPV-negative HNSCC tumors (Puram et al, 2018). Consistently, our analysis showed that SMYD3 expression negatively correlated with the expression of multiple type I IFN response and APM genes within individual cancer cells (**Fig.1G, Supplementary Figure 7**). These data suggest that SMYD3 is a key regulator of the expression of a broad repertoire of immune-related genes in HPV-negative HNSCC cells.

### SMYD3 mRNA expression is associated with repression of immune signatures in HPV-negative HNSCC human tumors

To determine whether the expression of SMYD3 is associated with enrichment of IFN-response and APM genes in human HPV-negative HNSCC tumor samples, we interrogated the Cancer Genome Atlas (TCGA) dataset of patients with primary HPV-negative HNSCC. Consistently with the aforementioned results, normalized enrichment scores of Hallmark gene sets revealed that higher SMYD3 mRNA expression correlated with suppression of immune signatures (**Fig. 1H**).

### UHRF1, a reader of H3K9me3, is downregulated after transient SMYD3 depletion and regulates the transcription of immune-related genes in HPV-negative HNSCC cells

SMYD3 is known to activate the transcription of downstream target genes, however, our data suggest a reverse phenotype, whereby SMYD3 is associated with repression of immune-related signatures. We thus reasoned that SMYD3 may regulate the expression of immune-related genes through an indirect mechanism, specifically through the upregulation of a repressive chromatin modifier or reader, which then in turn represses the expression of immune-related genes in HPV-negative HNSCC cells.

To identify candidate repressive epigenetic regulators, we curated and queried a list of 438 factors that are known to be involved in epigenetic regulation (**Supplementary Table 3**), we sorted the genes that had significantly decreased mRNA expression levels upon SMYD3 depletion and encoded for chromatin factors with known repressive functions on gene transcription. We found that the gene encoding the Ubiquitin Like PHD And Ring Finger Domain-containing protein 1 (UHRF1) had significantly lower mRNA expression after SMYD3 depletion (**Supplementary Figure 8**). UHRF1 is an E3 ubiquitin ligase which is necessary for the recruitment of DNMT1 to CpG DNA methylation sites and ensures high fidelity DNA maintenance methylation through DNMT1. UHRF1 binds and reads hemimethylated CpG marks through its SET- and RING-associated (SRA) domain, and H3K9me3 through its Tudor domain (TTD)-plant homeodomain (PHD), mediating transcriptional repression of its downstream target genes (Rothbart et al 2012, Cheng et al 2013, Karagianni et al, 2008, Xie et al, 2012). It is also associated with increased proliferation and poor prognosis in multiple cancer types (Kong et al, 2019). In order to validate UHRF1 as a downstream target of SMYD3, we performed RT-qPCR and Western blotting in two HPV-negative HNSCC cell lines (HN-6, HN-SCC-151) after siRNA-mediated SMYD3 depletion and confirmed significant downregulation of UHRF1 both at the mRNA and protein levels in the presence or absence of IFN-β (**Fig. 2A, B**).

**Figure 2.**
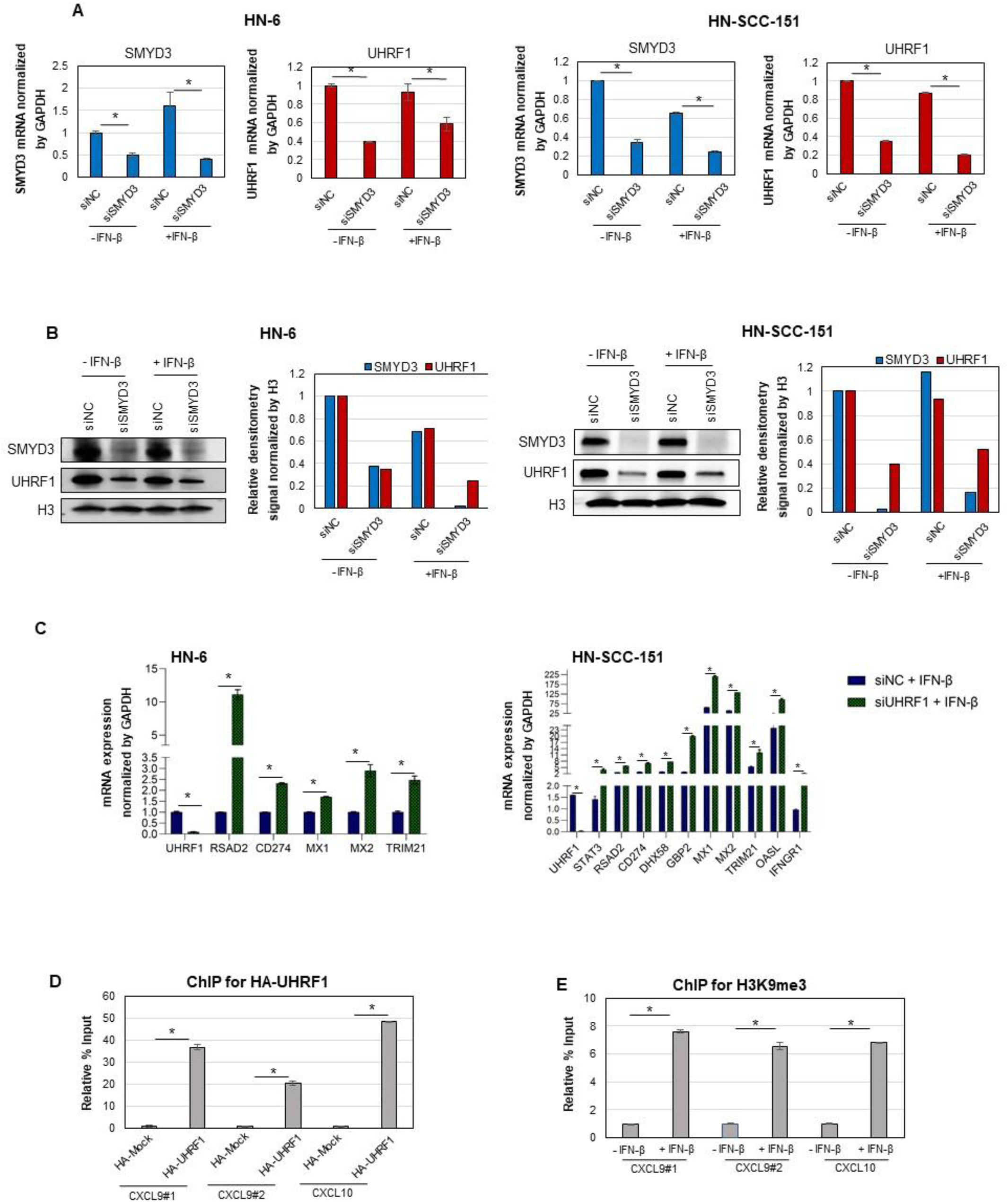
UHRF1 is transcriptionally regulated by *SMYD3* and silences the expression of immune-related genes in HPV-negative HNSCC cells through H3K9me3. **(A, B)** SMYD3 knockdown is associated with downregulation of UHRF1 in HN-6 and HN-SCC-151 cells at the mRNA **(A)** and protein levels **(B)**. **(C)** UHRF1 knockdown is associated with upregulation of immune-related genes in HN-6 and HN-SCC-151 cells. **(D)** UHRF1 binds on the promoters of two representative immune-related genes, CXCL9 and CXCL10. HN-6 cells were transfected with plasmids expressing HA-Mock or HA-UHRF1 for 48h and cells were exposed to IFN-β for 24h prior to collection. ChIP assays were conducted for HA followed by qPCR (SYBR green) for *CXCL9* and *CXCL10*. **(E)** The promoters of *CXCL9* and *CXCL10* are enriched for H3K9me3. ChIP assay for H3K9me3 in HN-6 cells before and after exposure of cells to INF-β.

To evaluate whether SMYD3 directly binds to and regulates the transcription of *UHRF1*, we conducted a ChIP assay targeting FLAG after transfection of HN-6 cells with FLAG-SMYD3 or FLAG-Mock expressing plasmids. We did not observe significant enrichment of FLAG-SMYD3 in the promoter region of *UHRF1*, suggesting that SMYD3 may be regulating the transcription of *UHRF1* indirectly (**Supplementary Figure 9**).

To assess whether UHRF1 regulates the expression of immune-related genes, HN-6 and HN- SCC-151 cells were transfected with either UHRF1-targeting (siUHRF1) or control siRNA (siNC) for 72 hours and treated with or without IFN-β for 24 hours. Similarly to SMYD3 depletion, UHRF1 depletion induced significant upregulation of multiple representative immune-related genes in both HNSCC cell lines, both in the presence (**Fig. 3C**) and absence of IFN-β exposure (**Supplementary Figure 10**).

**Figure 3.**
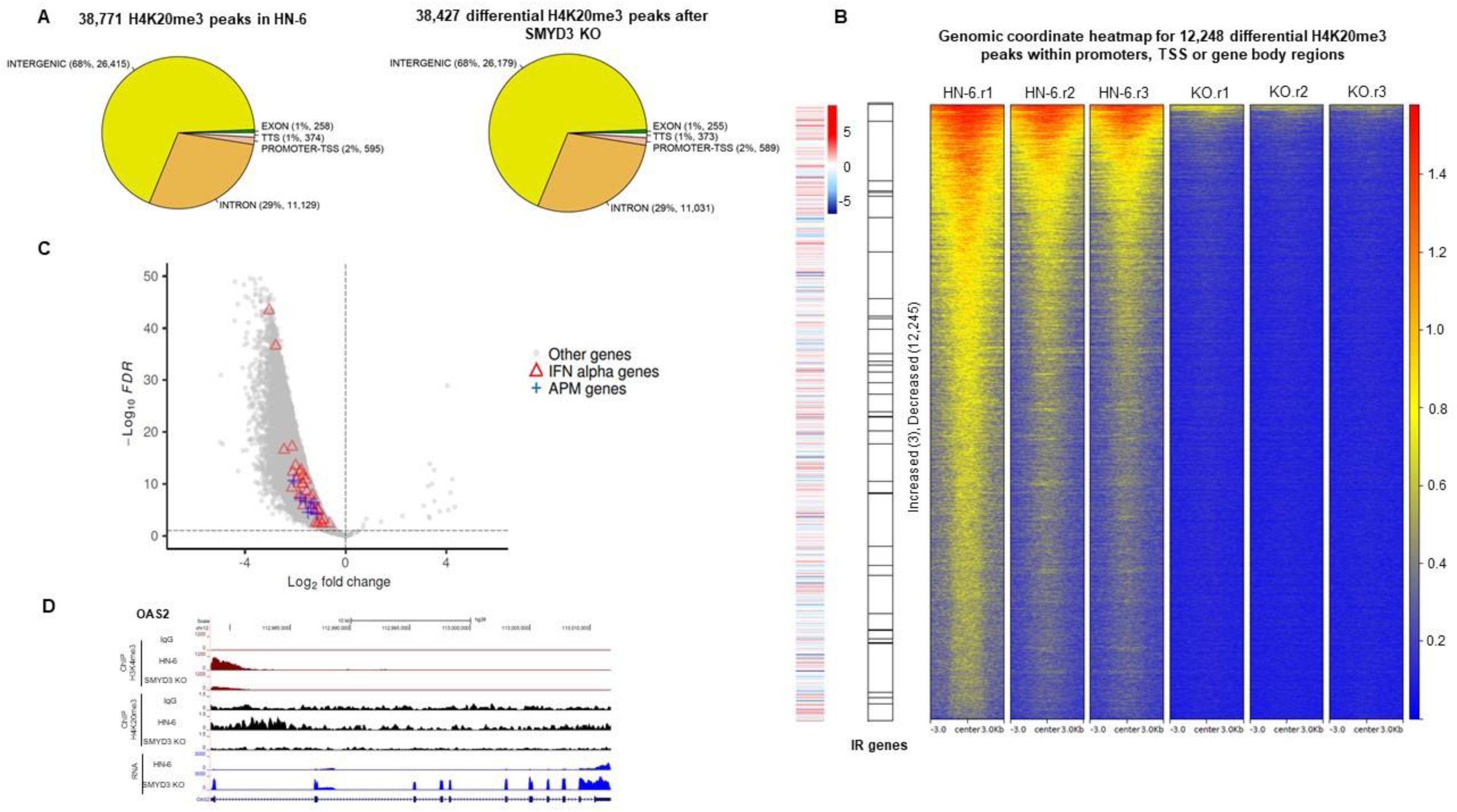
Permanent SMYD3 KO induces re-expression of immune-related genes through decreased deposition of H4K20me3. **(A)** Venn diagrams of peak distributions of H4K20me3 over genomic elements in HN-6 cells and after CRISPR-mediated SMYD3 KO (5-3 cell line). Cells were exposed for 24h to IFN-β. **(B)** Genomic coordinate heatmap of 12,248 H4K20me3 differential peaks present on promoters or gene body regions, altered in HN-6 cells after SMYD3 KO (5-3 cell line) and 24h of exposure to IFN-β. Peaks were aligned by the center of each peak. Only the gene coordinates overlapping with 12,248 differentially enriched H4K20me3 signals were used. Peaks were ordered from top to bottom based on Max tag density signals. Far left panel: concordant RNA-seq heatmap (log-fold change), middle panel: immune-related (IR) genes depicted as horizontal black lines in the black and white vertical column. **(C)** Volcano plot of DESeq2 results of H4K20me3 peaks (total = 38,784) in HN-6 cells and SMYD3 KO cells (5-3) after 24h of exposure to IFN-β. 38,408 H4K20me3 differential peaks were decreased and 19 H4K20me3 differential peaks were increased. FDR threshold: 0.1. Red triangles: 31 differential peaks overlapping with the promoters/TSS/gene bodies of 20 IFN alpha genes; blue crosses: 7 differential peaks overlapping with the promoters/TSS/gene bodies of 6 APM genes; gray circle: other genes. **(D)** UCSC browser tracks showing genome-wide mapping of H3K4me3 (maroon color), H4K20me3 (black color) and RNA-seq (blue color) of HN-6 cells or CRISPR SMYD3 KO cells after 24h of exposure to IFN-β. Tracks are focused on a representative immune-related gene, OAS2.

Given that UHRF1 represses its downstream genes, we hypothesized that UHRF1 may bind on the promoters of immune-related genes enriched with H3K9me3. To investigate whether UHRF1 directly binds and represses the expression of immune-related genes through H3K9me3, HN-6 cells were transfected with HA-UHRF1 or HA-Mock plasmids for 48h and exposed to IFN-β for 24h prior to cell collection. A ChIP-assay for the HA-tag followed by qPCR showed that HA-UHRF1 was enriched in the promoters of two representative immune-related genes, *CXCL9* and *CXCL10* (**Fig. 3D**), indicating a direct interaction between HA-UHRF1 and the promoters of these genes. Interestingly, a ChIP assay for H3K9me3 followed by qPCR for the same promoter sites of *CXCL9* and *CXCL10* showed enrichment of H3K9me3, which was significantly increased in the presence of IFN-β (**Fig. 3E**). We also attempted to conduct ChIP-seq for endogenously expressed human UHRF1 in HN-6 cells using a variety of commercially available antibodies, however we were not able to obtain a signal for UHRF1.

These results suggest that SMYD3 regulates the expression of immune-related genes through UHRF1, whereby UHRF1 binds to the promoters and represses the expression of immune-related genes enriched in H3K9me3.

### Permanent SMYD3 knockout decreases the occupancy of H4K20me3 in the promoters and gene body regions of immune-related genes in HPV-negative HNSCC cells

We then pursued to evaluate the protein expression levels of UHRF1 in SMYD3 KO cells. Interestingly, stable knockout of SMYD3 did not perturb steady-state UHRF1 levels, indicating that compensatory mechanisms restore UHRF1 expression after long-term SMYD3 loss (**Supplementary Figure 11**). We hypothesized that this may be related to the fact that UHRF1 depletion induced significant cell death in our HN-6 and HN-SCC-151 cell lines (data not shown), and that SMYD3 KO cell lines may compensate by upregulating UHRF1 expression levels back to a necessary minimum so that they can survive. This is in accordance with previous publications reporting that UHRF1 is necessary for the survival of a variety of cancer cell lines (Zhang et al, 2018, Li et al, 2019, Mudbhary et al, 2014).

Given that SMYD3 has been reported to write H4K20me3, a repressive mark, we hypothesized that SMYD3 regulates the deposition of H4K20me3 on immune-related genes, silencing their expression. To investigate this possibility further, we sought to evaluate the effect of SMYD3 KO on the genomic distribution of H4K20me3, and more particularly on the promoters and gene body regions of immune-related genes, using CUT&RUN assays in HN-6 and the CRISPR SMYD3 KO cell line 5-3 after IFN-β exposure. 38,771 H4K20me3 peaks were called in HN-6 cells. Of these peaks, 38,408 were significantly decreased, while 19 peaks were significantly increased in the SMYD3 KO cells (FDR<0.1) (**Fig.3A**). 68% of the differential peaks were distributed in intergenic regions and 29% in intronic regions, while less than 2% were distributed in promoters and TSS sites (**Fig. 3A**). 4,456 genes were annotated to 12,248 differential peaks overlapping with promoters, TSS or gene body regions, and of these genes, most of them (4,454 genes) were annotated to decreased H4K20me3 peaks (**Fig. 3B**). Among these 4,454 genes, 1,604 (36%) demonstrated significantly increased mRNA expression, signifying that SMYD3-mediated H4K20me3 is important in repressing these genes. 1,432 genes (32.2%) had decreased mRNA expression, and 1,418 genes (31.8%) showed no expression changes (FDR<0.1, absolute log2FC > 1.3) (**Fig.3B**).

Amongst the 4,456 genes that were annotated to promoters, TSS and gene body regions with differential H4K20me3 peaks, 26 immune-related genes were annotated to 38 decreased H4K20me3 peaks (**Fig.3C, Supplementary Table 4**). Of these, 20 genes were significantly upregulated (76.9%), 3 genes were unaltered (11.5%), and 3 genes were significantly downregulated (14.8%) (**Supplementary Fig.12**). The upregulation of the 20 genes out of 26 immune-related genes was found to be a statistically significant overrepresentation event (Fisher exact test, p<0.0001), indicating an immune-related gene specific upregulation by the loss of H4K20me3 marks. An example of H4K20me3 mapping on a representative immune-related gene, as well as respective RNA-seq tracks, are shown in **Fig. 3D**.

Interestingly, we also observed a global decrease in H3K4me3 peaks corresponding to immune-related genes in SMYD3 KO cells (**Supplementary Figure 13A, B**), which would have been expected to be associated with decreased transcription of immune-related genes. This finding suggests that H4K20me3 may predominate in regulating the transcriptional activation status of immune-related genes, even if H3K4me3, an activation mark, is depleted from these genes in SMYD3 KO cells. Furthermore, H3K9me3 peaks corresponding to immune-related genes were not affected with SMYD3 knockout (**Supplementary Figure 13C, D**), suggesting that SMYD3 does not affect the deposition of this mark on immune-related genes. This finding underscores that in the permanent SMYD3 KO state, the presence of H3K9me3 on immune-related genes may represent a complementary and SMYD3-independent mechanism of repression of immune-related genes.

These data suggest that SMYD3 silences the expression of immune-related genes through the deposition of H4K20me3 and that this repression may be reversed in a permanent SMYD3 KO system, inducing the upregulation of immune-related genes.

### SMYD3 depletion does not increase the expression of LINE retrotransposons in HPV-negative HNSCC cells

Given multiple reports supporting the importance of re-expression of transposable elements in the stimulation of type I IFN response in cancer cells (Chiappinelli et al 2015, Sheng et al, 2018, Kong et al, 2019), we sought to investigate whether SMYD3 affected the expression levels of LINEs. To evaluate this possibility, we conducted RNA-seq in HN-6 cells transfected with negative control or a SMYD3-targeting siRNA for 72h with IFN-β exposure. Results showed that the average mRNA expression levels of LINEs in control or SMYD3 depleted HN-6 cells were not significantly different, suggesting a LINE-independent role of SMYD3 in regulating type I IFN responses (**Supplementary Fig. 14**).

### Smyd3 depletion induces intratumoral CD8+ T-cell influx and enhances the antitumor efficacy of anti-PD-1 treatment in a syngeneic mouse model of HPV-negative HNSCC

Given that SMYD3 depletion led to transcriptional upregulation of multiple type I IFN response and APM genes in human HPV-negative HNSCC cell lines *in vitro*, we sought to evaluate the effect of Smyd3 depletion in the tumor immune microenvironment and its antitumor efficacy in an *in vivo* syngeneic mouse model of HPV-negative HNSCC cells. To this purpose, we utilized an established, DMBA (7,12-dimethylbenz(a)anthracene)-induced, HPV-negative mouse oral carcinoma 1 (MOC1) cell line in a syngeneic, heterotopic C57BL/6 mouse model of MOC1 flank tumors which is infiltrated with low numbers of CD8+ T-cells and is resistant to therapy with PD-1 inhibition, recapitulating immunotherapy-resistant HPV-negative HNSCC (Judd et al, 2012).

First, Smyd3 protein expression in MOC1 cells and the feasibility of knockdown of Smyd3 *in vitro* using Smyd3 ASOs were confirmed (**Supplementary Figure 15**). We then treated MOC1 tumors with Smyd3 or control ASOs with subcutaneous daily injections 5 days/week, and tumor volumes and weights were measured twice weekly. Two different doses were tested at 25mg/kg (control ASO n=4, Smyd3 ASO n=5) and 50mg/kg (control ASO n=5, Smyd3 ASO n=5), and mice were sacrificed on day 26 post-tumor cell implantation (day 20 of ASO treatment). No significant differences in the average tumor volumes were observed between control and Smyd3 ASO treated tumors (**Fig. 4A**, left top panel). Immunohistochemistry (IHC) for control or Smyd3 ASOs in MOC1 tumors demonstrated successful intratumoral penetration of the MOC1 tumor microenvironment by the ASOs (**Fig. 4A**, left bottom panel). Interestingly, CD8 IHC revealed significantly increased influx of CD8+ T-cells in MOC1 tumors treated with the 25mg/kg dose of Smyd3 ASOs compared to control ASO, while no significant difference was observed with the 50mg/kg dose (**Fig.5A**, right top and bottom panels). These results support that systemic treatment with Smyd3 ASOs increased the influx of intratumoral CD8+ T-cells in a syngeneic mouse model of MOC1 flank tumors, however, higher dosing seemed to have a deterring effect in intratumoral CD8+ T-cell trafficking. Furthermore, treatment with the lower dose of 25mg/kg of Smyd3 ASOs was associated with a lower average tumor volume of MOC1 tumors compared to control ASOs, however, this did not reach statistical significance.

**Figure 4.**
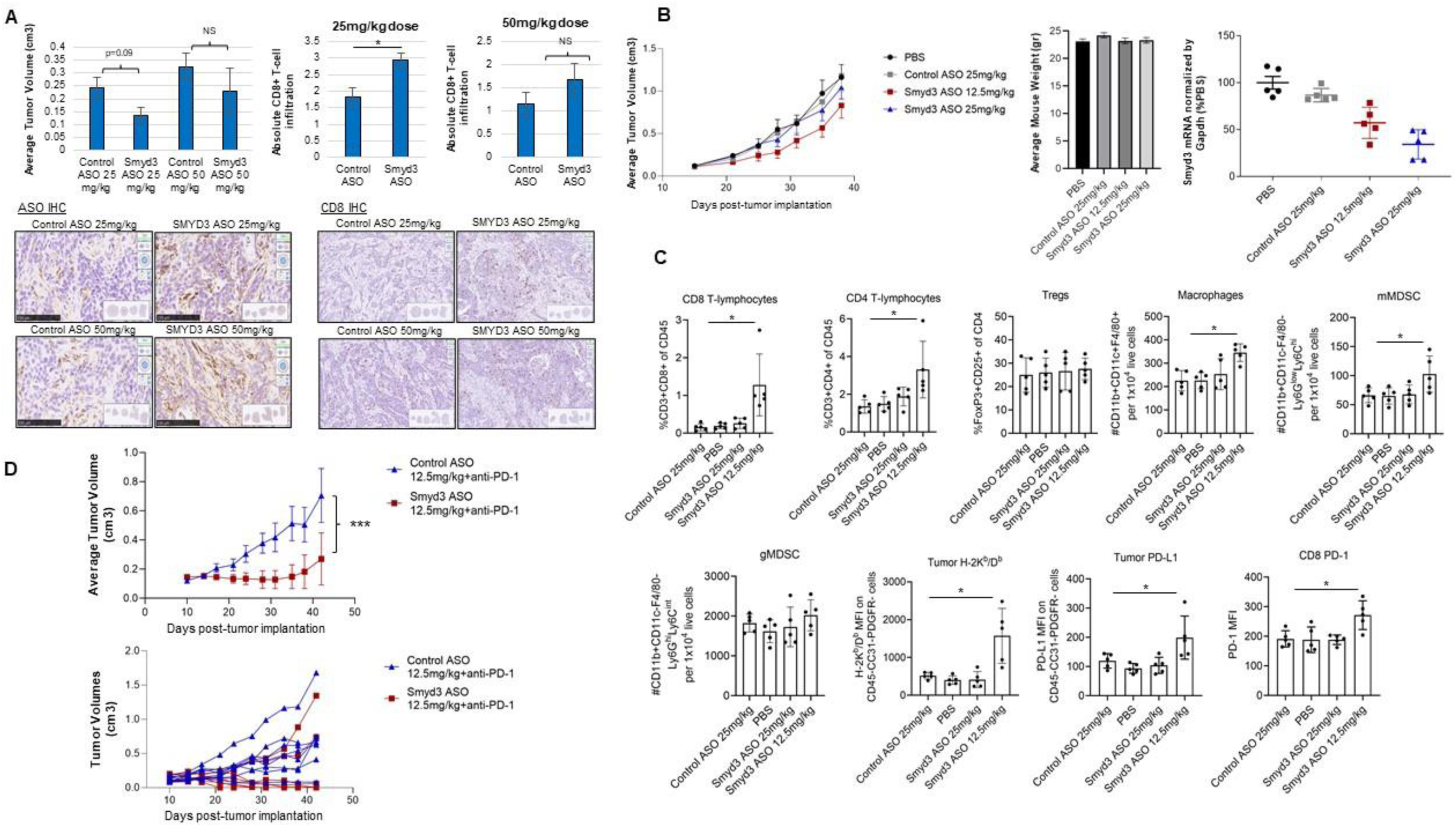
Smyd3 depletion increases CD8+ T-cell infiltration and induces significant tumor growth repression in combination with anti-PD-1 in a syngeneic mouse model of MOC1 tumors. **(A)** C57BL/6 mice were injected with 5×106 MOC1 cells in the right flank, and once they reached an average tumor volume of 0.05cm3 (day 6), treatment with Smyd3 or control ASOs was initiated at 25mg/kg or 50mg/kg with daily subcutaneous injections 5 days per week. On day 26 post-tumor implantation (day 20 of ASO treatment), mice were sacrificed, tumors were surgically resected and formalin fixed. Left top panel: average tumor volumes on day 26 post-tumor implantation, control ASO 25mg/kg: n=4, Smyd3 ASO 25mg/kg: n=5, control ASO 50mg/kg: n=5, Smyd3 ASO 50mg/kg: n=5. Data are mean +/- SEM, Student’s t-test at the indicated time point. Left bottom panel: Immunohistochemistry (IHC) for control or Smyd3 ASOs in MOC1 tumors at day 26 post-tumor cell implantation. Right top panel: Absolute CD8+ T-cell infiltration (CD8 IHC) in MOC1 tumors treated with control or Smyd3 ASOs at the indicated dosages (day 26 post-tumor cell implantation). Data are mean +/- SEM, Student’s t-test, *p-value=0.03, NS: non-significant. Right bottom panel: Representative examples of IHC for CD8 in MOC1 tumors treated with control or Smyd3 ASOs at the indicated dosages. **(B)** Left panel: Average tumor volumes of MOC1 tumors treated with PBS (n=10), control ASOs at 25mg/kg (n=10), Smyd3 ASOs at 25mg/kg (n=10) and 12.5mg/kg (n=10). Treatment with subcutaneous daily injections 5 days/week was initiated once tumors reached an average of 0.1cm3. Data are mean +/- SEM. Student’s t-tests between all group combinations were non-significant. Middle panel: Average weight of mice per treatment group (n=10 per group). Right panel: qPCR for Smyd3 mRNA of MOC1 tumors treated with PBS, control ASO 25mg/kg, Smyd3 ASO 12.5mg/kg and Smyd3 ASO 25mg/kg. RNA extraction was conducted from 5 tumors per group. Data are mean +/- SEM. **(C)** Multicolor flow cytometry of MOC1 tumors on day 39 post-tumor implantation (day 24 of treatment). MOC1 tumors treated with PBS, control ASOs at 25mg/kg, Smyd3 ASOs at 25mg/kg and at 12.5mg/kg were surgically resected and tumor cell digestion was conducted. Single-cell suspensions were incubated with antibodies conjugated to fluorophores and flow cytometry was conducted. Data are mean +/- SD (standard deviation). Student’s t-test between control ASO at 25mg/kg and Smyd3 ASO at 12.5mg/kg. *P<0.05, **P<0.01, ***P<0.001. **(D)** Top panel: Average tumor volumes of flank MOC1 tumors in C57BL/6 mice treated with control ASOs plus isotype IgG, Smyd3 ASOs plus isotype IgG, control ASOs plus anti-PD-1, Smyd3 ASOs plus anti-PD-1. Data are mean+/-SEM. Student’s t-test between compared groups. *P<0.05, **P<0.01, ***P<0.001. Bottom panel: Hairline growth curves of flank MOC1 tumors treated with control ASOs plus anti-PD-1 and Smyd3 ASOs plus anti-PD-1. This experiment was replicated with similar results one time. This experiment was replicated twice.

**Figure 5.**
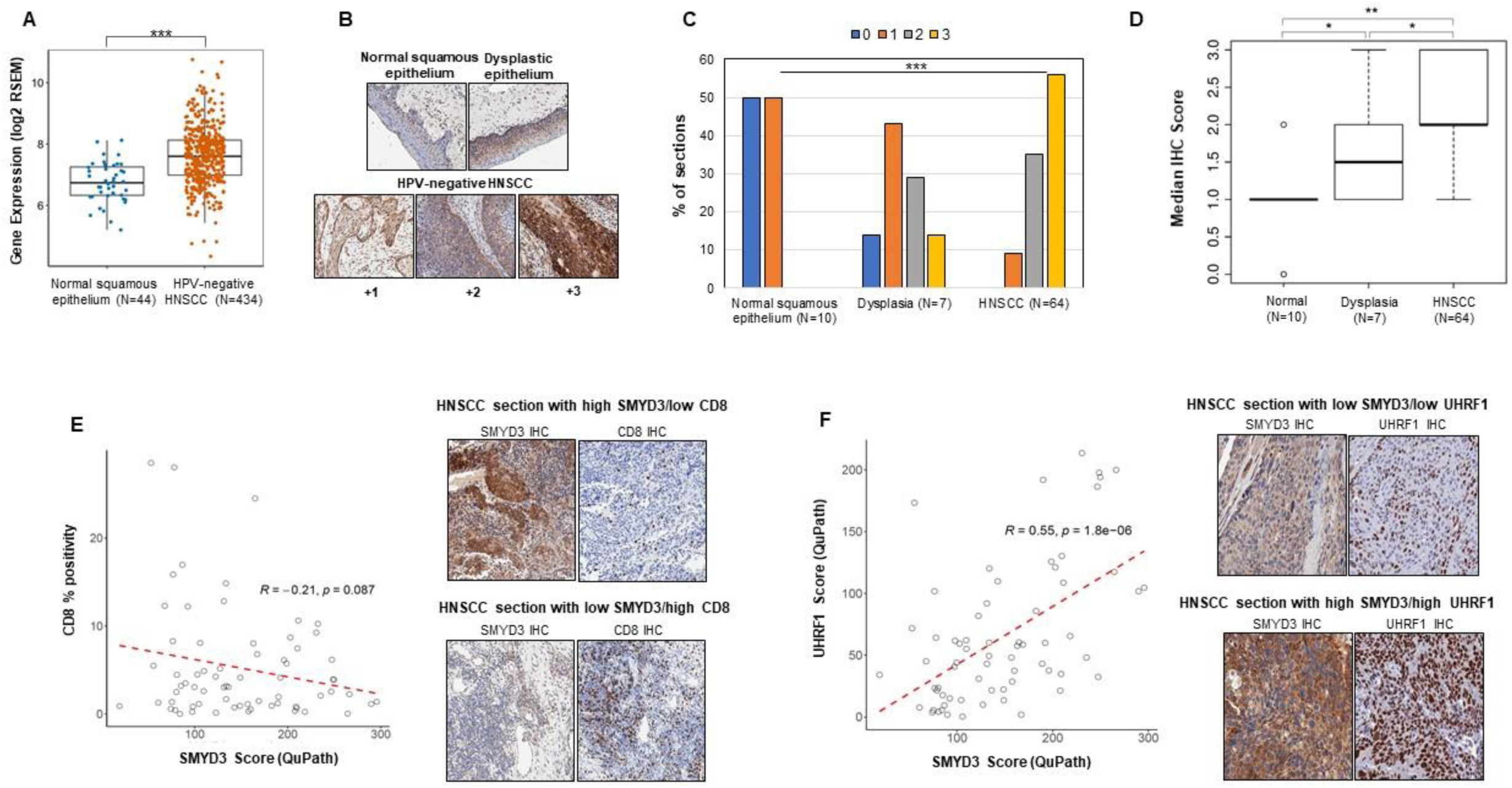
SMYD3 is overexpressed in HPV-negative HNSCC tumors and is associated with CD8+ T-cell infiltration and UHRF1 protein expression. **(A)** SMYD3 mRNA expression levels are higher in HPV-negative HNSCC tumors compared to normal squamous epithelium. TCGA database analysis of 434 HPV-negative tumors and 43 normal squamous epithelial samples. ***P<0.001. **(B)** Representative images of immunohistochemical (IHC) staining for SMYD3 in normal squamous epithelium, dysplastic epithelium and HPV-negative HNSCC sections. **(C)** Distribution of IHC scores among normal squamous epithelium (N=10), dysplastic epithelium (N=7) and HPV-negative SCCHN tumors (N=64). Kruskal Wallis test; *** P<0.001. **(D)** Median IHC score for SMYD3 in normal squamous epithelium (N=10), dysplastic epithelium (N=7) and HPV-negative HNSCC tumors (N=64). Dunn test for multiple comparisons; * p<0.05, ** p <0.001. **(E)** Left panel: Correlation of SMYD3 and CD8 immunohistochemical staining in 64 HPV-negative HNSCC tumors; QuPath scores, Person coefficient r=-0.21, p=0.087. Right panel: Representative images of IHC staining of an HPV-negative HNSCC tumor section with high SMYD3/low CD8 staining, and an HNSCC section with low SMYD3/high CD8 staining. **(F)** Left panel: Correlation of SMYD3 and UHRF1 immunohistochemical staining in 64 HPV-negative HNSCC tumors; QuPath scores, Person coefficient r=0.55, p<0.001. Right panel: Representative images of IHC staining of an HPV-negative HNSCC tumor section with high SMYD3/high UHRF1 staining, and a HNSCC section with low SMYD3/low UHRF1 staining.

To validate these observations and to further explore the effect of Smyd3 ASOs in the tumor microenvironment, we conducted multicolor flow cytometry of MOC1 tumors treated with PBS (n=10), control ASO at 25mg/kg (n=10), Smyd3 ASOs at 25mg/kg (n=10) and Smyd3 ASOs at 12.5mg/kg (n=10). We opted to treat mice at an even lower dose of Smyd3 ASOs (12.5mg/kg), considering that 25mg/kg induced greater influx of CD8+ T-cells compared to 50mg/kg; we thus reasoned that an even lower dose could be associated with an even greater influx of CD8+ T-cells. As per the aforementioned experiment, no significant differences were observed between the treatment groups, although MOC1 tumors treated with Smyd3 ASOs at 12.5mg/kg tended to have smaller average tumor volumes at all time points compared to the control ASO treated tumors (**Fig.4B**, left panel). The treatment with ASOs was tolerated well at all doses (**Fig.4B**, middle panel) and mice were sacrificed on day 39 post-tumor cell implantation (day 24 of treatment). MOC1 tumors were surgically resected, and tumor RNA-extraction and multicolor flow were conducted (n=5 tumors per group). Smyd3 mRNA levels from tumor extracts were decreased in a dose-dependent manner, with approximately a 50% decrease and a 75% decrease observed in tumors treated with Smyd3 ASOs at 12.5mg/kg and 25mg/kg respectively (**Fig.4B**, right panel). CD8+ and CD4+ T-cells were significantly increased in the MOC1 tumors treated with Smyd3 ASOs at 12.5mg/kg compared to PBS, control ASO and Smyd3 ASO treated tumors at 25mg/kg (**Fig. 4C**). Macrophages were also significantly increased in the tumors treated with the Smyd3 ASOs at 12.5mg/kg compared to all other groups. Tregs and granulocytic MDSCs were not increased, however an influx of monocytic MDSCs was observed in the tumors treated with the Smyd3 ASOs at 12.5mg/kg. Consistently with the aforementioned RNA-seq data in human HPV-negative HNSCC cell lines, MHC class I H-2K^b^ and PD-L1 were also significantly upregulated on MOC1 tumor cells. Furthermore, PD-1 expression was significantly upregulated on CD8+ T-cells, suggesting a state of exhaustion.

We then sought to evaluate whether Smyd3 ASOs could synergize with anti-PD-1 therapy in this syngeneic mouse model of flank MOC1 tumors. MOC1 tumors were implanted in the right flanks of C57BL/6 mice, and once the average tumor volume reached 0.1cm^3^, treatment was initiated with control ASOs at 12.5mg/kg plus anti-PD-1 (200ug) (n=8) and Smyd3 ASOs at 12.5mg/kg plus anti-PD-1 (200ug). Mice were injected subcutaneously with the ASOs for 5 days/week and intraperitoneally twice weekly with anti-PD-1. The group treated with the Smyd3 ASO plus anti-PD-1 combination demonstrated a statistically significant decrease in the average MOC1 tumor volumes compared to control ASO plus anti-PD-1 (**Fig. 4D**). 4/8 mice treated with the Smyd3 ASO plus anti-PD-1 combination were cured, while 2/8 mice had significant tumor regressions (0.06cm^3^ and 0.007cm^3^ at the time of sacrifice). On the other hand, the MOC1 tumors in 2/8 mice of the Smyd3 ASO combination group “escaped” the treatment effect and grew significantly, similarly to the control ASO plus anti-PD-1 treated group.

The above data support that Smyd3 depletion induces favorable changes in the MOC1 tumor microenvironment with influx of CD8+, CD4+ T-cells and macrophages, while also upregulating the expression of H-2K^b^ and PD-L1 on the cell surface of MOC1 tumors cells. These changes are associated with sensitization of MOC1 tumors to anti-PD-1 therapy, with deep responses and complete tumor regressions in mice treated with systemic Smyd3 ASOs plus anti-PD-1 therapy.

### SMYD3 is overexpressed in human HPV-negative HNSCC tumors and is associated with CD8+ T-cell infiltration and UHRF1 protein expression

To assess the expression pattern of SMYD3 in HPV-negative HNSCC tumors, the HPV-negative HNSCC database of the TCGA was interrogated and results showed that SMYD3 mRNA levels were significantly higher in HPV-negative HNSCC tumors compared to normal squamous epithelium (**Fig. 5A**). To validate these results at the protein level, we conducted immunohistochemistry (IHC) for SMYD3 in 64 HPV-negative tumor samples, as well as in 10 available normal and 7 dysplastic buccal squamous epithelium samples (**Fig. 5B**). Analysis of the staining results using a semiquantitative scoring scale (0, +1, +2, +3 staining intensity) revealed that the percentage of samples with +3 SMYD3 staining increased significantly from normal, to dysplastic epithelium and then to squamous cell carcinoma samples (**Fig. 5C**). Furthermore, approximately 80% of tumors showed strong staining (+2, +3). Concordantly, the median IHC score was significantly higher in dysplastic samples compared to normal squamous samples (Dunn test for multiple comparisons, p<0.05), as well as between squamous carcinoma and dysplastic samples (Dunn test for multiple comparisons, p<0.05) and squamous carcinoma and normal squamous samples (Dunn test for multiple comparisons, p<0.01) (**Fig. 5D**). These data support a potential role of SMYD3 in the oncogenesis of HPV-negative HNSCC.

Furthermore, we sought to evaluate the relative expression levels of SMYD3 in the cancer cell compartment compared to the stroma. Using the QuPath software, the average H-score of the cancer to the stroma compartment in sections of HPV-negative HNSCC tumors was significantly higher (Wilcoxon rank sum test, p=3.6×10^-6^) (**Supplementary Figure 16A**). This finding was further corroborated by the significantly higher mRNA expression levels of SMYD3 in HPV-negative HNSCC cancer cells of a publicly available single-cell RNA-seq database (Puram et al, 2018) compared to other cell subtypes, such as T-cells, B-cells, dendritic cells, macrophages and fibroblasts (Wilcoxon rank sum test, p<0.0001) (**Supplementary Figure 16B**). These data support a more prominent role of SMYD3 in regulating biological functions in cancer cells and less in stroma/immune cells, however, this needs to be more definitively answered through experimental interrogation of the role of SMYD3 in immune cell function.

In our previously published work (Vougiouklakis et al, 2017), we showed that higher SMYD3 mRNA expression levels were associated with significantly lower levels of CD8+T-cell attracting chemokines, such as CXCL9 and CXCL10, as well as lower CD8A mRNA levels in HPV-negative HNSCC tumors (TCGA). Based on this as well the aforementioned preclinical data supporting that SMYD3 depletion induces re-expression of immune-related genes and influx of CD8+ T-cells in mouse MOC1 tumors, we pursued to evaluate the association between SMYD3 protein levels and CD8 T-cell infiltration in HPV-negative HNSCC tumors. To this purpose, we conducted IHC for CD8 in the aforementioned cohort of 64 newly diagnosed HPV-negative HNSCC patients. As expected, we found that tumors with higher SMYD3 protein expression levels tended to have lower CD8+ T-cell infiltration (R=-0.21, p=0.087, **Fig. 5E**).

Given our finding that UHRF1 is a downstream target of SMYD3, we sought to evaluate whether SMYD3 protein expression levels correlate with UHRF1 protein levels in HPV-negative HNSCC tumors. Indeed, quantification of SMYD3 and UHRF1 protein expression levels in the aforementioned HPV-negative HNSCC tumor sections showed that SMYD3 correlated positively with UHRF1 (Pearson’s coefficient R=0.55, p<0.001) (**Fig. 5F**). We also investigated the correlation between UHRF1 protein levels and CD8 infiltration, however, we did not find a statistically significant correlation (R=-0.011, p=0.93, **Supplementary Figure 17**).

### SMYD3 protein levels predict pathologic response to neoadjuvant pembrolizumab

We then sought to evaluate whether SMYD3 expression levels predict response to treatment with pembrolizumab. To this purpose, we used a published RNA-seq database of 20 evaluable patients with newly diagnosed, treatment naïve HPV-negative oral cavity HNSCC who were treated with one dose of neoadjuvant pembrolizumab and then were surgically resected 2-3 weeks after the treatment to evaluate pathologic tumor response (PTR) and clinical to pathological downstaging (Uppaluri et al, 2020). Higher SMYD3 mRNA levels at baseline were significantly associated with no clinical to pathological downstaging (Wilcoxon rank sum test, p=0.02, **Fig.6A**, left graph). While higher baseline UHRF1 mRNA expression levels were also observed in patients without clinical to pathological downstaging, this difference did not reach statistical significance (Wilcoxon rank sum test, p=0.49, **Fig.6A**, middle graph). Similarly and as expected, lower CD8 mRNA levels were also observed in patients with no clinical to pathological downstaging, however did not reach statistical significance (Wilcoxon rank sum test, p=0.09, **Fig.6A**, right graph).

**Figure 6.**
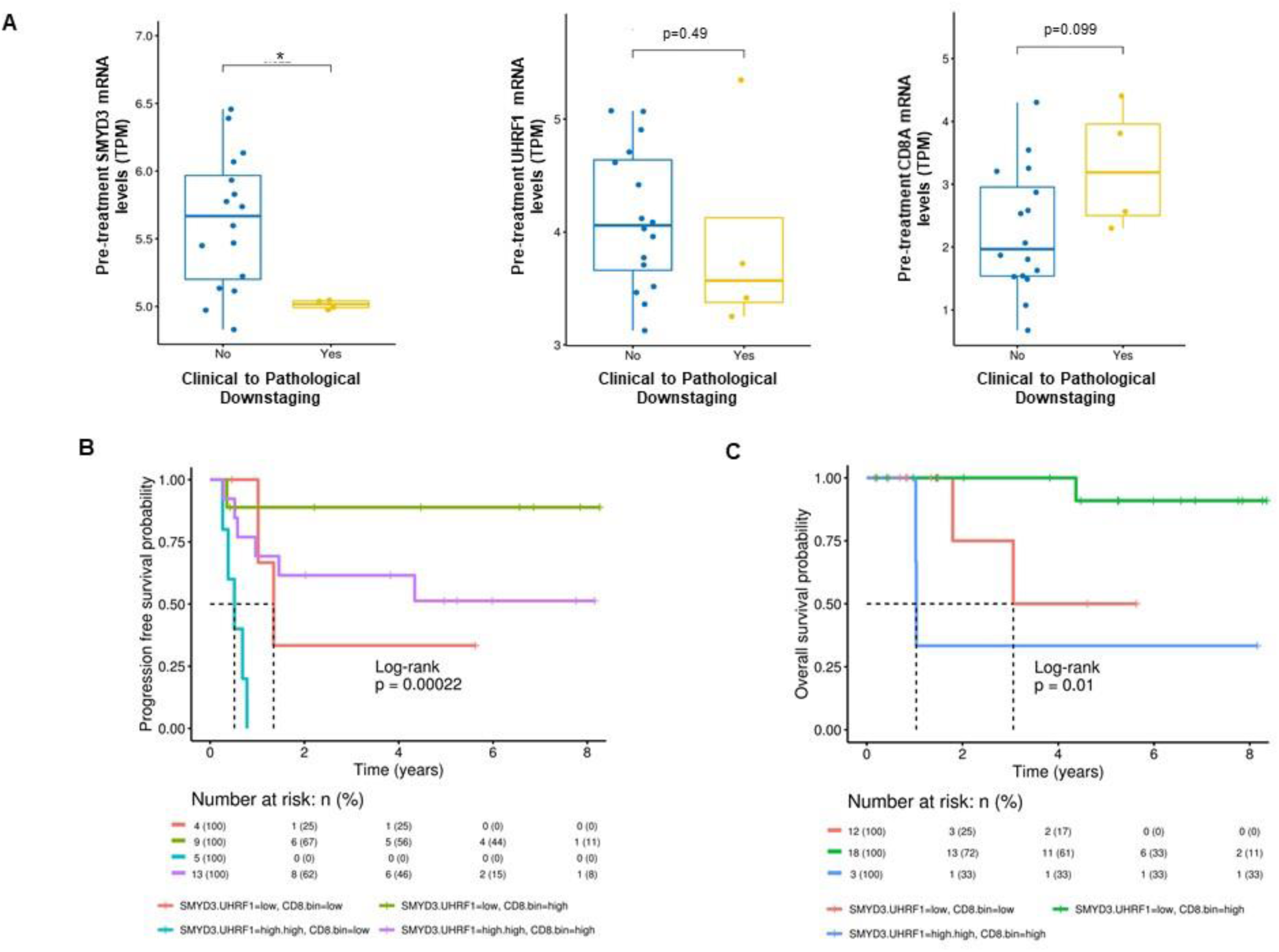
SMYD3 expression predicts response to neoadjuvant pembrolizumab and determines survival when combined with UHRF1 and CD8 expression. **(A)** Boxplot showing the correlation between pretreatment SMYD3 (left panel), UHRF1 (middle panel) and CD8 (right panel) mRNA expression levels and clinical to pathological downstaging after one dose of neoadjuvant pembrolizumab in 20 oral cavity, treatment-naïve HPV-negative HNSCC patients. N=16 patients with no clinical to pathological downstaging, N=4 patients with clinical to pathological downstaging. Wilcoxon rank sum test for SMYD3, p=0.022, for UHRF1, p=0.49, for CD8, p=0.09. **(B)** Kaplan Meier progression free survival (PFS) curve in 31 HPV-negative HSNCC patients treated with standard of care surgery and/or chemoradiotherapy. Patients with higher SMYD3/UHRF1 and low CD8 protein expression levels at baseline had significantly worse PFS compared to patients with low SMYD3/UHRF1 and high CD8 baseline protein expression levels. Log-rank p-0.0002. **(C)** Kaplan Meier overall survival (OS) curve in 33 HPV-negative HSNCC patients treated with standard of care surgery and/or chemoradiotherapy. Patients with higher SMYD3/UHRF1 and low CD8 protein expression levels at baseline had significantly worse OS compared to patients with low SMYD3/UHRF1 and high CD8 baseline protein expression levels. Log-rank p-0.01.

### Combined SMYD3, UHRF1 and CD8 expression predict progression free and overall survival in patients with primary HPV-negative HNSCC

We then assessed whether baseline SMYD3 and UHRF1 protein expression levels correlate with progression free (PFS) and overall survival (OS). Interestingly, while SMYD3 or UHRF1 baseline protein expression levels alone were not significantly associated with PFS or OS in our cohort of 35 evaluable patients (**Supplementary Figure 18**), combined protein expression of SMYD3, UHRF1 and CD8 at baseline predicted a significant impact on both PFS and OS. More specifically, patients with high baseline SMYD3/UHRF1 and low CD8 protein expression had significantly worse OS and PFS compared to patients with low baseline SMYD3/UHRF1 and high CD8 protein expression levels (**Fig. 6B, C**).

## DISCUSSION

Although checkpoint inhibition with pembrolizumab or nivolumab has established a new standard of care therapeutic paradigm for HPV-negative HNSCC, the majority of these patients do not respond to this therapy. Deciphering mechanisms that drive checkpoint inhibitor resistance in HPV-negative HNSCC is thus of paramount importance.

In the present study, we demonstrate that genetic depletion of SMYD3 by siRNAs, ASOs or CRISPR increases cancer cell sensitivity to IFN-β by transcriptionally derepressing the expression and leading to an orchestrated induction of multiple type I IFN response and APM genes in HPV-negative HNSCC cells. Transcriptional repression of these genes is dependent on SMYD3-mediated transcriptional regulation of Ubiquitin-Like PHD And RING Finger Domain-Containing Protein 1 (UHRF1), a key epigenetic regulator that reads the repressive trimethylated lysine 9 on histone H3 (H3K9me3) mark, and on the deposition of H4K20me3 on immune-related genes. Smyd3 depletion with systemic Smyd3 ASO treatment in an anti-PD-1 resistant immunocompetent mouse model of flank MOC1 tumors induces influx of CD8+ T-cells, upregulates mouse PD-L1 and MHC class I molecules, and increases sensitivity to anti-PD-1 therapy. Finally, SMYD3 is overexpressed in HPV-negative HNSCC tumors and baseline SMYD3 and UHRF1 mRNA tumor expression levels predict response to anti-PD-1 neoadjuvant therapy in HPV-negative HNSCC patients.

Our work reveals that depletion of SMYD3, a H3K4 and H4K20 methyltransferase, induces derepression of type I IFN response and APM genes in HPV-negative HNSCC cell lines through two temporally distinct, chromatin-based mechanisms (**Figure 7**). Transient SMYD3 knockdown induced downregulation of UHRF1, an epigenetic reader of H3K9me3, while both UHRF1 and H3K9me3 were found enriched in the promoters of prototypical type I IFN response genes *CXCL9* and *CXCL10*, suggesting that UHRF1 may bind on the promoters of these genes and silence their expression. Genome-wide mapping of UHRF1 could help solidify the role of UHRF1 in directly regulating the transcriptional repression of immune-related genes. Furthermore, it would be important to decipher the mechanisms of enrichment of H3K9me3 on the promoters of *CXCL9* and *CXCL10* genes, as reversal of this repressive mark may have synergistic or additive effects to SMYD3 depletion towards derepression of type I IFN response and APM genes. Additionally, identifying proteins that interact with UHRF1 and specifically direct it to the promoters of immune-related genes would also be important to further understand the mechanism through which UHRF1 may silence these genes.

**Figure 7.**
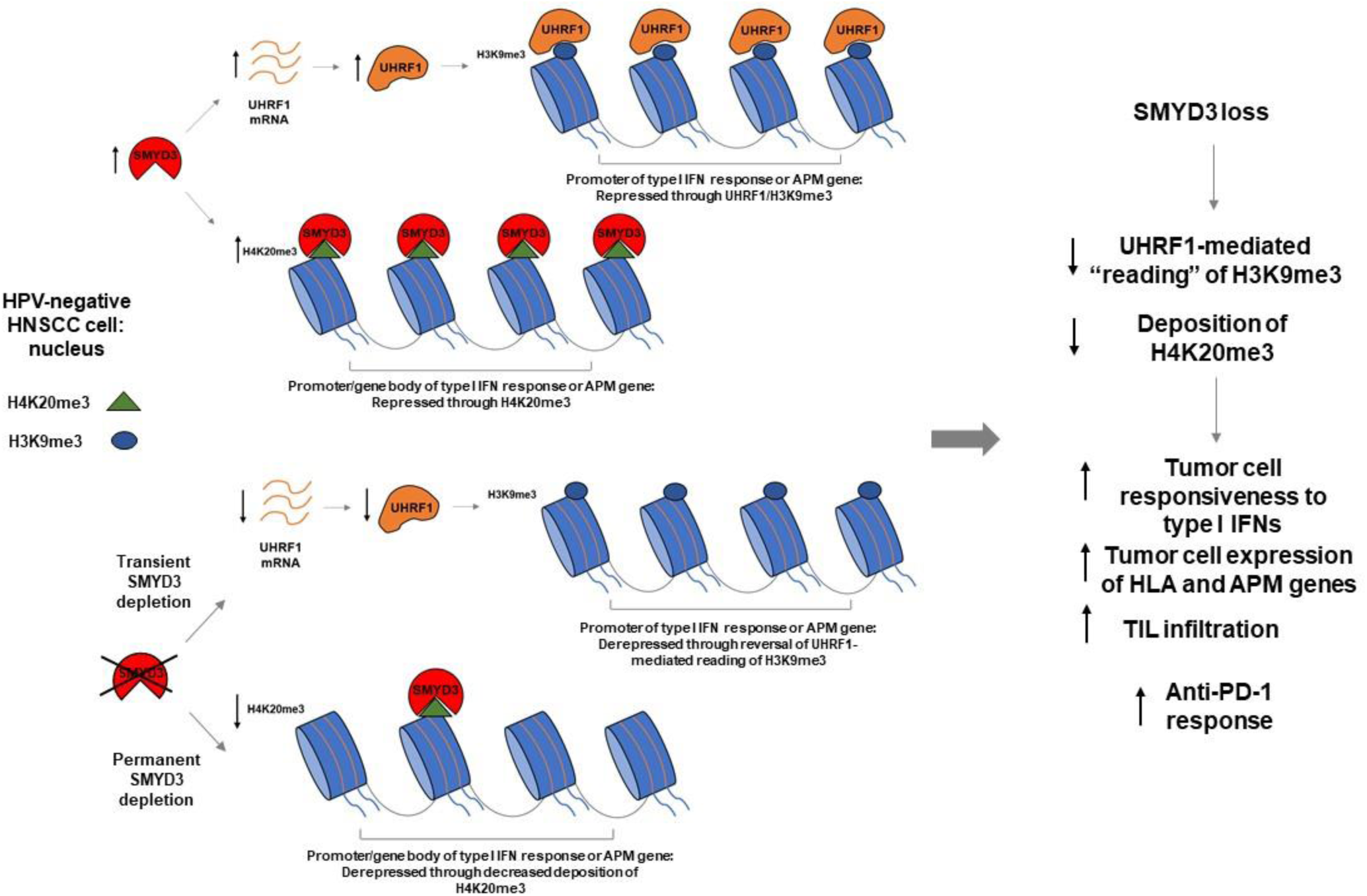
Schematic of the mechanism of SMYD3-mediated repression of immune-related genes.

Permanent SMYD3 knockout decreased the deposition of H4K20me3, a repressive mark, on intragenic regions of type I IFN response and APM genes in our CRISPR SMYD3 knockout cell lines. While a massive decrease in the deposition of H4K20me3 was observed with SMYD3 depletion in the majority of genes corresponding to H4K20me3 peaks, a significantly higher percentage of immune-related genes (69%) were transcriptionally activated compared to random genes (42%), implying specificity of the repressive effect of SMYD3 through H4K20me3 for immune-related genes. While we observed downregulation of UHRF1 at the mRNA and protein level after siRNA-mediated SMYD3 knockdown, we could not validate this in the permanently SMYD3 depleted CRISPR knockout HNSCC cell lines, which maintained UHRF1 protein expression levels. We hypothesize that this may be related to the fact that UHRF1 depletion caused significant cell death in our HPV-negative HNSCC cell lines, suggesting that its expression is necessary for the survival of these cells. Despite the maintained expression of UHRF1 in our SMYD3 CRISPR knockout cell lines, upregulation of type I IFN response and APM genes was maintained as a phenotype. Our data suggest that this was mediated through depletion of H4K20me3 from intragenic regions of immune-related genes, which may be the predominant mechanism of immunomodulation in permanently SMYD3 depleted cells. Genome-wide mapping for SMYD3 to assess whether it directly binds and co-occupies immune-related genes with H4K20me3 will be important to further solidify the proposed mechanism of action of SMYD3 as an immunomodulator in HNSCC cells.

The described mechanisms are distinct and complement evidence regarding the functions of certain epigenetic modifiers as regulators of antitumor immunity in different cancer types (Cao et al, 2020). Well studied mechanisms that have been reported pertain to the repression of endogenous retroviral elements (ERVs) or dsRNAs, type I IFN response and APM genes through DNA methylation (DNA methyltransferases), H3K27 trimethylation mediated by the protein methyltransferase Enhancer of zeste homologue 2 (EZH2), and H3K9 trimethylation by SETDB1 in melanoma or non-small cell lung cancer cell lines (Peng et al, 2015, Nagarsheth et al, 2016, Chiappinelli et al, 2015, Topper et al, 2017, Griffin et al, 2021). More specifically, recent studies have demonstrated activation of type I interferon responses through re-expression of dsRNAs and repetitive elements, such as ERVs, induced by inhibition of DNA methylation alone or together with histone deacetylase (HDAC) inhibitors in mouse melanoma models (Chiappinelli et al, 2015, Topper et al 2017). Furthermore, SETDB1 amplification has been associated with immune exclusion and resistance to PD-1 inhibition, while SETDB1 loss derepresses transposable elements (TEs), immunostimulatory genes and TE-encoded retroviral antigens through erasure of H3K9me3, inducing TE-specific cytotoxic CD8+ T-cell antitumor immune responses in melanoma and Lewis lung carcinoma mouse models (Griffin et al, 2021). Additionally, lysine-specific histone demethylase 1 (LSD1) has also been reported to repress the expression of ERVs and increase the stability of certain RNA-induced silencing complex (RISC) components, ultimately leading to reduced expression of ERVs and thus to diminution of intracellular type I IFN responses in breast cancer cells (Sheng et al, 2018). Depletion of LSD1 increases the expression of ERVs and activates type I interferon responses, stimulating tumor immunogenicity through upregulation of major histocompatibility complex (MHC) class I expression and inducing T-cell infiltration and anti-tumor immunity in mouse melanoma models (Sheng et al, 2018).

Given multiple studies highlighting the importance of the re-expression of retransposons and ERVs as a mechanism of induction of type I IFN response by cancer cells (Chiappinelli et al, 2015, Sheng et al, 2018, Griffin et al, 2021), we also sought to evaluate whether SMYD3 may affect the expression of some of these elements in HPV-negative HNSCC cells. Our RNA-seq database of HN-6 cells treated with siSMYD3 did not reveal any expression changes in the retransposon sequences (LINE) with SMYD3 knockdown (**Supplementary Figure 14**), however, this analysis will need to be further supplemented with total RNA-seq to evaluate changes in the expression levels of ERVs. Concordantly with the lack of expression changes of retransposons, the *Stimulator of Interferon Response CGAMP Interactor 1* (*STING1*) gene expression levels, which are expected to be upregulated with dsRNA intracellular stress, were not upregulated in our siSMYD3 and CRISPR SMYD3 knockout RNA-seq databases (**Supplementary Figure 19**). Other studies have demonstrated the importance of EZH2-mediated deposition of H3K27me3 directly on key Th1-type chemokine genes *CXCL9* and *CXCL10*. More specifically, EZH2 and DNA-methyltransferase 1 (DNMT-1) transcriptionally repress Th1-type chemokine genes *CXCL9* and *CXCL10*, and inhibition of both EZH2 and DNMT-1 increases CD8+ T-cell trafficking, reduces tumor growth and improves the efficacy of PD-1 blockade in mouse ovarian cancer models (Peng et al 2015). Similarly, EZH2 transcriptionally represses the expression of *CXCL9* and *CXCL10* through H3K27 trimethylation in colon cancer cells, and overexpression of EZH2 is associated with decreased CD8+ and CD4+ T-cell tumor infiltration and poor prognosis in colon cancer patients (Nagarsheth et al, 2016). EZH2-mediated deposition of H3K27me3 has also been demonstrated on the *B2M* promoter in MOC1 cells, and EZH2 inhibition enhances antigen presentation and responses to anti-PD-1 immunotherapy in the anti-PD-1 MOC1 resistant mouse model of HPV-negative HNSCC (Zhou et al, 2019). An important question to decipher is whether different repressive histone marks, such as H3K27me3 or H4K20me3, have mutually exclusive, complementary or redundant functions in the repression of immune-related genes, thus allowing for synergistic roles or the development of resistance mechanisms through the respective epigenetic regulators governing their deposition or erasure.

Using Smyd3 ASOs, mouse Smyd3 depletion induced marked influx of CD8+ T-cells, CD4+ T-cells and macrophages, and markedly upregulated the expression of MHC class H-2K^b^ on MOC1 tumors, inducing sensitization of the anti-PD-1 resistant HPV-negative MOC1 mouse model to anti-PD-1 therapy. Tregs and granulocytic MDSCs, which have also been shown to play a significant role as immunosuppressive immune cell subsets in HPV-negative HNSCC (Greene et al, 2020), were not increased with Smyd3 ASO treatment. However, treatment with Smyd3 ASOs induced upregulation of PD-1 and PD-L1, indicating an immune checkpoint-mediated exhaustive state of the CD8+ T-cells, which may explain why monotherapy with Smyd3 ASOs did not induce any significant tumor growth restraint despite the marked influx of CD8+ T-cells. In this setting, combination treatment of MOC1 tumors with Smyd3 ASOs and anti-PD-1 led to cure of 4/8 mice and marked tumor growth restraint in 2/8 mice (**Fig.4D**). Smyd3 ASO treatment was also associated with influx of monocytic MDSCs, which may explain why 2/8 treated mice escaped the treatment effect of the Smyd3 ASOs and anti-PD-1 combination. Further interrogation of possible mechanisms of escape is warranted and ongoing, with a focus on potential competing and redundant functions of other chromatin modifiers that could orchestrate the deposition of repressive histone marks on immune-related genes. Another important observation is that a higher dose of Smyd3 ASOs did not induce influx of CD8 and CD4+ T-cells, suggesting a deleterious effect of higher Smyd3 ASO dosage on the intratumoral trafficking of T-cells. As such, the elucidation of the differential effect of Smyd3 depletion on cancer versus immune cells would be of paramount importance.

60-80% of HPV-negative HNSCC tumor sections in our cohort of 64 patients overexpressed SMYD3, and increased baseline SMYD3 protein expression in HPV-negative HNSCC tumor samples predicted poor pathologic response to neoadjuvant pembrolizumab, supporting that SMYD3 may abrogate therapeutic responses to pembrolizumab. Furthermore, high baseline SMYD3 protein levels in combination with high UHRF1 and low CD8 levels predicted poor OS and PFS in HPV-negative HNSCC patients treated with standard of care chemoradiotherapy. The fact that effective antitumor immunity is necessary for the efficacy of chemoradiotherapy in HNSCC explains our finding that SMYD3 expression, when combined with UHRF1 and CD8 baseline expression levels, affects the survival outcomes of HNSCC patients treated with chemoradiotherapy. While these results are conducive to the hypothesis that SMYD3 diminishes the IFN responsiveness and antigen presentation capacity of HPV-negative HNSCC cancer cells, our patient cohort sample was small and further validation would be needed in a larger cohort of patients.

Interestingly, studies have suggested that certain highly inducible inflammatory genes are maintained in transcriptional repression by nuclear repressor complexes, such as the nuclear repressor coreceptor 1 (NCoR), the corepressor of REST (CoREST) or the glucocorticoid receptor (GR) (Ghisletti et al., 2009; Hargreaves et al., 2009; Hoberg et al., 2006; Huang et al., 2009; 2011; Ogawa et al., 2004; Pascual et al., 2005; Saijo et al., 2009; Flammer and Rogatsky, 2011). Upon appropriate pro-inflammatory stimuli, these transcriptional repressor complexes are removed and the inflammatory genes transition from their basal, silenced state to an actively transcribed state. Along these lines, Stender et al (2012) previously reported the importance of H4K20 methylation/demethylation in the transcriptional regulation of certain TLR4-responsive inflammatory genes through SMYD5, which trimethylates H4K20, and PHF2, a H4K20 demethylase which is activated upon lipopolysaccharide (LPS)-induced TLR4 stimulation. SMYD5 was found to be a component of the repressor NCoR complex and importantly, knockdown of the other four members of the SMYD family in mouse macrophages did not induce upregulation of TLR4-responsive inflammatory genes, indicating distinct biological roles for each SMYD family member. Furthermore, SMYD5 knockdown also did not alter the inhibitory effect of the GR receptor on inflammatory genes, signifying a pathway-specific mechanism of repression of these genes. In another study (Xu et al, 2015), Smyd2 was found to repress the expression of *Il-6* and *Tnf* through H3K36 dimethylation of their promoters, inhibiting macrophage activation and M1 polarization, and to increase the expression of *Tgf-β* which promoted Treg differentiation. Nagata et al (2015) also reported that Smyd3 induces the expression of *Foxp3* through a Tgf-β/Smad3 dependent mechanism and promotes Treg differentiation, and that *Smyd3* knockout mice demonstrated excessive RSV-induced pulmonary inflammation and damage secondary to uncontrolled inflammatory responses within the lungs of *Smyd3* knockout mice. These data suggest that the SMYD family members may have distinct biological functions in the regulation and optimization of anti-viral and anti-bacterial immunity.

Our study provides important insight into how epigenetic mechanisms mediated by SMYD3 may silence anti-tumor immunity and drive resistance to checkpoint inhibition by anti-PD-1 therapy in HPV-negative HNSCC. In summary, SMYD3 depletion increases sensitivity of HPV-negative HNSCC cancer cells to IFN-β by inducing derepression of type I IFN response and APM genes. This occurs through downregulation of UHRF1, abrogating the “reading” of H3K9me3 which is enriched on the promoters of immune-related genes in transiently SMYD3-depleted cells, and through erasure of the repressive H4K20me3 mark on the gene bodies of immune-related genes in permanently SMYD3 depleted cells. This induces favorable changes in the tumor microenvironment, with influx of CD8+, CD4+ T-cells and macrophages, marked upregulation of MHC class I on the surface of tumor cells, and ultimately leads to sensitization to anti-PD-1 therapy in an anti-PD-1 resistant syngeneic HPV-negative HNSCC mouse model. As a therapeutic targeting strategy for SMYD3, ASOs provide a promising therapeutic platform and could overcome the significant problem of specificity when targeting methyltransferase enzymes. Furthermore, ASOs have the potential to abolish oncogenic mechanisms induced both by enzymatic hyperactivity due to mutations as well as overexpression of the target gene. The latter is relevant for SMYD3 which is overexpressed but rarely mutated (<2%) in HPV-negative HNSCC (**Supplementary Figure 20**). Two ASOs have been FDA approved for the treatment of hypercholesterolemia and spinal muscular atrophy, further underscoring the translational feasibility and promise of this drug platform. These data support a rational translational strategy, whereby HPV-negative HNSCC patients could be stratified by baseline SMYD3 protein expression levels and treated with combinatorial SMYD3 ASOs with PD-1 checkpoint inhibition to increase clinical responses to checkpoint immunotherapy.

## ACKNOWLEDGEMENTS

This work was funded by the Intramural Research Program of the National Cancer Institute.

## AUTHOR CONTRIBUTIONS

NN: Data curation, investigation, formal analysis, methodology, validation, visualization, writing-original draft; BB: Data curation, investigation, formal analysis, methodology, validation, visualization, writing-original draft, KS: conceptualization, data curation, formal analysis, investigation, methodology, software, visualization, writing-original draft; KB: data curation, investigation, methodology, validation, visualization; DT: conceptualization, data curation, investigation, methodology, visualization; YR: data curation, formal analysis, investigation, methodology; CS: conceptualization, data curation, formal analysis, investigation, methodology, visualization, writing-review and editing; AC: conceptualization, investigation, methodology; RLB: data curation, investigation, methodology; TTT: data curation, investigation, methodology, validation, visualization; BC: data curation, methodology, writing-review and editing; RB: data curation, formal analysis, methodology; LR: data curation, formal analysis, investigation, methodology, writing-review and editing; MWL: data curation, investigation; HS: data curation, formal analysis, investigation, methodology; EFE: data curation, formal analysis, investigation, methodology; HC: data curation, formal analysis, investigation, methodology; XL: : data curation, formal analysis, investigation, methodology; BK: : data curation, formal analysis, investigation, methodology, visualization; CC: data curation; MM: data curation; SS: data curation, investigation, methodology; KT: data curation, investigation, methodology; YN: conceptualization; RU: data curation, investigation, methodology; JBS: methodology, investigation; CVW: funding acquisition, writing-review and editing; JDL conceptualization, investigation, methodology; GLH; funding acquisition, conceptualization, investigation, methodology; VS: conceptualization, data curation, formal analysis, funding acquisition, investigation, methodology, project supervision, visualization, writing-original draft, review and editing.

## INCLUSION AND DIVERSITY

We worked to ensure gender balance in the inclusion of human subjects.

## RESOURCE AVAILABILITY

### Lead Contact

Further information and requests for resources and reagents should be directed and will be fulfilled by the Lead Contact, Vassiliki Saloura.

### Materials availability

This study did not generate new unique reagents.

### Data availability

Previously unpublished datasets used and/or generated during this study will become publicly available if the manuscript is favorably recommended for peer review.

## EXPERIMENTAL MODEL AND SUBJECT DETAILS

All murine experiments were performed using 4-6 week-old female C57BL/6 mice from Taconic. All animals were maintained in accordance with the NCI-Bethesda Animal Care and Use Committee.

## METHOD DETAILS

### Cell culture

HPV-negative squamous cell carcinoma cell lines HN-6 and HN-SCC-151 were derived from patients with locoregionally advanced HNSCC and were kindly provided by Dr. Tanguy Seiwert (University of Chicago). HN-6 cells were maintained in DMEM medium with 10% fetal bovine serum, 1% penicillin/streptomycin, and 2 nM L-glutamine. HN-SCC-151 cells were maintained in DMEM/F12 medium, 10% fetal bovine serum, 1% penicillin/streptomycin and 2 nM L-glutamine.

### Generation of *SMYD3* knockout cell lines using CRISPR

SMYD3 CRISPR knockout cell lines (*SMYD3* KO 5-2, 5-3) were generated from parental HN-6 cells using clustered regularly interspaced short palindromic repeats (CRISPR/Cas9) technology. After single-cell selection of GFP-expressing HN-6 cells using flow cytometry, SMYD3 expression levels were assessed by Western blotting to confirm efficiency of knockout (**Supplementary Fig.21**).

### siRNA transfections

siRNA oligonucleotides were purchased from Millipore-Sigma to target the human SMYD3 mRNA (SASI_Hs02_0035-5988) and UHRF1 mRNA (SASI_Hs02_00311672). For convention, SMYD3-targeting siRNA is referred to as siSMYD3 and UHRF1-targeting siRNA is referred to as siUHRF1. The negative control siRNA was purchased from Dharmacon (siRNA negative control Dharmacon ON-TARGET plus control pool, #D-001810-10-20, Horizon Discovery, Lafayette, CO). HNSCC cells were plated overnight in 10cm dishes and were transfected with siRNA duplexes (50 nM final concentration) using Lipofectamine RNAimax (Thermo Fisher Scientific, Grand Island, NY) for 72h (3 days) or 144 h (6 days). For 6 days of siRNA transfection, re-transfections were performed on day 3.

### Interferon-β treatment

For all experiments, interferon-β treatments were performed at a concentration of 1000U/mL for 24 hours (R&D systems Inc., Minneapolis, MN). When siRNA transfections were performed for a 72 hour total duration, interferon-β treatment was initiated at the 48 hour time point and lasted for 24 hours prior to cell collection.

### Western blotting

Nuclear extracts were prepared using the Nuclear Complex Co-IP kit (Active Motif) and 10 μg of each extract was loaded to examine protein levels of SMYD3, UHRF1 and Histone 3. Primary antibodies used were anti-SMYD3 (ab187149, Abcam, Cambridge, MA, dilution 1:2000), anti- UHRF1 (D6G8E, Cell Signaling Technologies, Danvers, MA, dilution 1:1000), and anti-histone 3 (ab1791, Abcam, Cambridge, MA, dilution 1:20000). For detection of histone mark H3K4me3, nuclear extracts were prepared using the Nuclear Complex Co-IP kit (Active Motif); 2 μg of each extract was loaded for each experiment and histone H3 was examined as loading control. The anti-H3K4me3 antibody was used (ab8580, Abcam, Cambridge, MA, dilution 1:1000). Densitometry of all the western blots was performed using ImageJ software (NIH, Bethesda, MD).

### Quantitative real-time PCR

Primers for human *GAPDH* (housekeeping gene), *SMYD3, UHRF1, TRIM21, STAT1, OASL, IFNGR1, OAS3, CD274, OAS2, DHX58, NCOA7, MX1, MX2, GBP2, RSAD2, TAP2, CANX, CXCL9,* and *CXCL10* were purchased from Sigma-Aldrich. RNA extraction was performed using the Zymo Research Direct-zol RNA miniprep kit (Zymo Research, Irvine, CA). cDNA conversion was performed using the Invitrogen SuperScript III First-Strand Synthesis System (Invitrogen, Carlsbad, CA). PCR was conducted in technical triplicates using either SYBR Select Master Mix (Applied Biosystems, Foster City, CA) or GreenLink qPCR Master-Mix (BioLink Laboratories, Washington D.C.). Each master mix was validated and showed similar results. PCR reactions were performed using CFX96 Touch Real-Time PCR Detection System (Bio-Rad Laboratories, Hercules, CA).

### ChIP-qPCR

ChIP-qPCR was performed on HN-6 cells for H3K9me3, HA-tagged UHRF1 and FLAG-tagged. For ChIP of H3K9me3, HN-6 cells were plated overnight at 40% confluence in 15cm dishes and were exposed to human interferon-β on next day for 24h. For the FLAG-SMYD3 and HA-UHRF1 ChIP, 15cm dishes of 40% confluence of HN-6 cells were transfected for 48h with plasmids for FLAG-MOCK or FLAG-SMYD3, or plasmids for HA-MOCK or HA-UHRF1, using Fugene HD. Cells were exposed to human interferon-β for 24h prior to harvesting cells. Cells were then trypsinized, resuspended and cross-linked with 1% formaldehyde for 5-8 min. Fixed cells were washed twice with 1X PBS, lysed using ChIP Lysis Buffer and sonicated using Bioruptor (Diagenode) as described above. The supernatant following sonication was diluted 5-fold using ChIP dilution buffer, and 50ul was used as input and the remaining for immunoprecipitation. The diluted supernatant was precleared with Invitrogen Dynabeads Protein G (Invitrogen 10004D) for 1h at 4oC and incubated with antibodies for H3K9me3 (Ab8898, Abcam, Cambridge, MA), FLAG (F3165, Sigma, St. Louis, MO) or HA (3724S, Cell Signaling Technology, Danvers, MA) overnight. On the next day, samples were incubated with prewashed Invitrogen Dynabeads Protein G beads for 2h at 4C, washed and reverse cross-linked overnight. All samples were purified using phenol-chloroform precipitation, resuspended in H2O and quantified using Qubit. Purified ChIP DNA was tested for human ChIP primers specific to the promoter regions of *UHRF1, CXCL9,* and *CXCL10*. For the *UHRF1* promoter region (FLAG-SMYD3 ChIP), primer sets consisting of the 153 bp region (nucleotides −166 to −14; sense: 5-GGCTGTACAGGAGGACTGGA-3 and antisense: 5-AGCAAAAACCCCCATCAGTT-3) or the *UHRF1* distal promoter region (nucleotides −3217 to −3123; sense: 5-CTCCCAAAGTGCTGGGATTA-3, and antisense: 5-GG CAACAAGAGCAAAACTCC-3) were used. For the *CXCL9* promoter region, a primer set close to the TSS (sense: 5’-TGCACTCCAATCAGAACCAG-3’, and antisense: 5’-CCAATACAGGAGTGACTTGGAAC-3’), and the region upstream of the TSS at -1000bp (sense: 5’-CGGTGTGATACCACCTTACAC-3’, anti-sense: 5’-GTTCCCTGATCACCAAGTCC-3’) were used. For the *CXCL10* promoter, a primer set close to the TSS sense: 5’-TCCCTCCCTAATTCTGATTGG-3’, and antisense: 5’-AGCAGAGGGAAATTCCGTAAC-3’) was used. PCR was performed in technical duplicates using GreenLink qPCR Master-Mix. PCR as a percent of the input was calculated for each ChIP sample using the formula: 100∗ 2^[CtInput – log2(IP volume/Input volume)-Ct IP], where IP volume/Input volume is the input dilution factor. Ct values of 2 technical replicates and at 2 biological replicates were used for the analysis.

### RNA-seq

RNA-seq was performed in HN-6 cells treated with SMYD3 targeting siRNAs, SMYD3 ASOs and in the CRISPR SMYD3 knockout cell lines 5-2 and 5-3. For the siRNA mediated SMYD3 knockdown, HN-6 cells were plated overnight at 40% confluence and transfected with siNC or siSMYD3 using Lipofectamine RNAimax (Thermo Fisher Scientific, Grand Island, NY) for 72h, with or without exposure to human IFN-β for 24h. For the SMYD3 ASO treatment, 40% confluent HN-6 cells were treated with SMYD3 ASOs or PBS for 72h, with or without exposure to human IFN-β for 24h. For the CRISPR SMYD3 knockout cell lines, parental HN-6 cells and the CRISPR SMYD3 knockout cell lines 5-2 and 5-3 were plated at 60% confluence, and collected after exposure or not to human IFN-β for 24h. Following completion of incubation, cells were trypsinized, washed twice with PBS, centrifuged and processed for RNA extraction (Direct-zol RNA miniprep kit, Zymo Research, Irvine, CA). Three biological replicates for each sample were processed to extract RNA, quantified using Qubit and sequenced. Samples were pooled and sequenced on NovaSeqStandard_SP using Illumina TruSeq Stranded mRNA Library Prep and paired-end sequencing. The samples have 68 to 130 million pass filter reads. Reads of the samples were trimmed for adapters and low-quality bases using Cutadapt before alignment with the reference genome (hg38) and the annotated transcripts using STAR. The samples had 62-75% non-duplicate reads. In addition, the gene expression quantification analysis was performed for all samples using STAR/RSEM tools. The raw counts are provided as part of the data delivery.

### CUT&RUN assays and DNA-sequencing

For CUT&RUN assays, the 14-1048 CUT&RUN kit by EpiCypher was utilized according to EpiCypher’s protocol. Briefly, CUTANA spike-in dNuc controls (H3K4me0, 1, 2, 3) were mixed together with washed streptavidin (SA) beads in 4 separate 1.5ml tubes, and incubated for 30min at RT on nutator. Concanavalin (ConA) beads were activated using cold bead activation buffer, washed twice using a magnet, resuspended in cold activation buffer, added at 10uL/sample in separate strip tubes (1 tube per experimental sample) and kept on ice. 500,000 cells per experimental condition were obtained after trypsinization from respective cell culture dishes (1 dish per biological replicate) and were washed with PBS x 3 to remove excess trypsin. Cells were then resuspended in 100uL/sample of RT wash buffer and washed twice at 600xg for 3min. After the final wash, cell pellets were resuspended in 105uL of RT wash buffer, and 100uL per sample were aliquoted into each 8-strip tube containing 10uL of activated beads. The cell-bead slurries were incubated on the benchtop for 10min at RT to allow for adsorption of the cells to the beads. After the incubation, the slurries were placed on a magnet and a small aliquot of the supernatant was obtained to confirm adsorption of cells to the beads (binding efficacy >93% of cell input). The supernatants were completed removed and the cell-bead slurries were then exposed to cold antibody buffer and vortexed. The CUTANA H3K4MetSTat spike-in control dNucs were added to designated positive (H3K4me3) and negative (IgG) control tubes. Then, 0.5ug of antibodies to H4K20me3 (ThermoFisher, MA5-36090), H3K4me3 (EpiCypher, 13-0041) or H3K9me3 (ab176916-GR246504-14) were added to each designated experimental tube. Biological triplicates were used for each experimental condition. The samples were incubated overnight on a nutator at 4oC. Next day, the 8-strip tubes containing the samples were placed on a magnet till the slurries cleared, supernatants were removed and the cell-beads were washed twice with cold cell permealization buffer. After the final wash, 50uL of the cold cell permealization buffer was added to the cell-bead slurries, and then 2.5uL of pAG-MNase was added to each sample. Samples were incubated for 10min at RT and the 8-strip tubes were placed back on a magnet. Supernatants were removed and cell-beads complexes were washed twice with cold cell permealization buffer. After the final wash, 50uL of cell cell permealization buffer was added in each sample, and targeted chromatin digestion followed by adding 1uL of chromatin digest additive to each sample. Strips were incubated for 2h at 4oC on a nutator and the reaction was stopped using Stop buffer. 0.5ng of spike-in Ecoli DNA was added to each sample and samples were incubated for 10min at 37oC in a thermocycler. The strips were then placed on a magnet and the supernatants containing the CUT&RUN enriched DNA were transferred to new tubes. DNA was purified per EpiCypher’s protocol, and library construction was conducted. Nucleic acid size selection to enrich for fragment sizes between 200-500bp was conducted using SPRIselect (Beckman Coulter Life Sciences, B23318). Samples were pooled and sequenced on NextSeq2000 using TruSeq ChIP and Swift Bioscience Accel-NGS 2S Plus DNA Library Prep Kits and paired-end sequencing. All the samples had yields between 53 and 80 million pass filter reads. Samples were trimmed for adapters using Cutadapt before the alignment. The trimmed reads were aligned with hg38 reference using Bowtie2 alignment. All the samples had library complexity with percent non-duplicated reads ranging from 75 to 88%.

### Immunohistochemistry

Tissue microarrays containing clinically annotated patient tumor samples were obtained from the Human Tissue Research Center of the University of Chicago Pathology Department (IRB#8980). 64 HPV-negative HSNCC tumors, 10 dysplastic lesions and 10 samples from normal buccal epithelium which stained for SMYD3 (abcam 187149), CD8 (DAKO, Cat#M7103, Clone: C8/144B) and UHRF1 (ab194236) using immunohistochemistry (IHC). For the SMYD3 IHC, the staining was performed on Leica Bond RX automated stainer. After deparaffinization and rehydration, tissue sections were treated with antigen retrieval solution (Leica Microsystems, AR9640) with heat near 100°C for 20 minutes. The anti-SMYD3 antibody (1:400) was applied on tissue sections for 1 hour incubation at room temperature. The antigen-antibody binding was detected with Leica Bond Polymer Refine Detection (Leica, DS9800) system and the slides were covered with cover glasses. For the CD8 IHC, the slides were stained on Leica Bond RX automatic stainer using the protocol “HTRC Bond DAB Refine”. Epitope retrieval solution I (Leica Biosystems, AR9961) was used for 20 minutes. The anti-CD8 antibody (1:400) was applied on tissue sections for 25 minutes incubation and the antigen-antibody binding was detected with Bond polymer refine detection (Leica Biosystems, DS9800). The tissue sections were covered with cover glasses. For the UHRF1 IHC, after deparaffinization and rehydration, tissue sections were treated with antigen retrieval buffer (S2367, DAKO) in a steamer for 20 minutes. The anti-UHRF1 antibody (1:200) was applied on tissue sections for 1 hour incubation at room temperature in a humidity chamber. Following TBS wash, the antigen-antibody binding was detected by Bond Polymer Refine Detection (DS9800, Leica Microsystems) and DAB+ chromogen (DAKO, K3468). Tissue sections were briefly immersed in hematoxylin for counterstaining and were covered with cover glasses. The IHC staining was approved by the Institutional Review Board of the University of Chicago (IRB#18-0468-AM002).

The stained tissue microarray slides were then scanned and digital image analysis algorithms were developed in QuPath v.0.3.0 (Belfast, UK) (Bankhead P et al, 2017). Digital slides were reviewed for 64 head and neck cancer tumors and included H&E stained sections along with IHC on serial sections for SMYD3, UHRF1 and CD8. All tissue sections had minimal artifact and staining pattern were consistent with specific staining. For each case, tumor was annotated to exclude normal tissue, artifact and necrotic regions. For SMYD3 and UHRF1 assessment, only tumor compartments were annotated from each core, while for the CD8 assessment, both the tumor and stroma compartments were evaluated together in each core. Parameters for cell detection were optimized and validated. SMYD3 immunolabeling was commonly positive in cancer cells, most commonly with a cytoplasmic pattern but with a detectable nuclear pattern too, while UHRF1 immunolabeling was predominantly nuclear. SMYD3 and UHRF1 immunolabeling were reported with an H-score. SMYD3 H-scores ranged from 84.14-295.65, while UHRF1 H-scores ranged from 0.3-213.53. CD8 scoring was reported as CD8% in each tumor core and ranged from 0.03-20.66%.

For the IHC staining of ASOs, slides were stained with rabbit polyclonal ASO (Ionis) antibody on a Ventana Ultra staining system. ASO slides were treated enzymatically with trypsin (Sigma, T8003). The slides were then blocked with endogenous biotin blocking kit (Ventana, 760-050) and normal goat serum (Jackson Immuno Labs, 005-000-121). The primary antibody was diluted with discovery antibody diluent (Ventana, 760-108) and incubated for 1 hour at 37°c. The antibodies were detected with biotin labeled goat anti-rabbit secondary antibody (Jackson Immuno Labs, 111-005-003). The secondary antibody was labeled with DABMap Kit (Ventana, 760-124). Images were scanned on a Hamamatsu S360 scanner at 20X resolution.

### Mice and in vivo mouse experiments

All animal experimental protocols were approved by the NCI-Bethesda Animal Care and Use Committee. 4-6 week-old female C57BL/6 mice were purchased from Taconic and used for the described experiments. The study designs and animal usage were conducted accordingly to all applicable guidelines by the NCI-Bethesda Animal Care and Use Committee. MOC1 cells were grown in vitro and were inoculated by subcutaneous injections of 5 million MOC1 cells in suspension using Matrigel, in the right flanks of C57BL/6 mice. Once flank tumors reached an average volume of 0.1cm3, mice were randomized into treatment groups and treatment was initiated according to each experiment. Mice were treated with control ASOs or Smyd3 ASOs at concentrations ranging from 12.5m/kg to 50mg/kg or PBS, as described in each experiment, for 5 days per week. Intraperitoneal injections of isotype IgG (InVivoMAb rat IgG2a, 2A3, 1mg/ml) or anti-PD-1 (InVivo Plus anti-mouse PD-1, RMPI-14) were conducted at 200ug/injection, twice weekly. Tumor length (L) and width (W) were measured twice weekly with calipers and tumor volumes were calculated using the formula LxW^2/2. Weights were measured twice weekly.

### Multicolor flow cytometry

For the multicolor flow of MOC1 tumors, mice were euthanized and flank MOC1 tumors were surgically resected, and mechanically and chemically digested into single-cell suspensions using the gentleMACS Dissociator and the mouse tumor dissociation kit by Miltenyi Biotec (130-096-730) respectively, per manufacturer’s protocol. Single-cell suspensions were filtered through 70uμ filters and washed with 1% BSA in PBS. Samples were incubated with anti-CD16/32 (Biolegend) antibody to block nonspecific staining. Subsequently, the primary antibodies were added and incubation for 30min was carried out in the dark. Cell surface staining was performed using fluorophore-conjugated anti-mouse CD45.2 (clone 104), CD3 (145-2C11), CD4 (GK1.5), CD8 (53-6.7), CD31 (390), PDGFR (APA5), PD-L1 (10F.9G2), H2-K^b^ (AF6-88.5), PD-1 (RMP1-30), CD11b (M1/70), Ly6G (1A8), Ly6C (HK1.4), CD11c (N418) and F4/80 (BM8) from Biolegend.

FoxP3+ regulatory T-cell staining was performed with the mouse regulatory T-Cell Staining Kit #1 (eBioscience) as per manufacturer’s protocol. Cell viability was assessed with Sytox (Thermo) or Zombie (Biolegend) dyes. All analyses were performed on a BD Fortessa analyzer running FACSDiva software and interpreted using FlowJo V.X10.0.7r2.

## QUANTIFICATION AND STATISTICAL ANALYSIS

### Statistical Analyses and Software

#### Lists of type I IFN response and APM genes

Based on previous publications (Chiappinelli et al, 2015, Sheng et al, 2018), we comprised a list of type I IFN response and APM genes (**Supplementary Table 1**). Bioinformatic interrogation of this gene list is shown in Figure 1. Genes that were not found to be expressed at the mRNA level in the RNA-seq databases of each cell system presented in Figures 1A-C were omitted. We also interrogated the gene set from the HALLMARK_INTERFERON_ALPHA_RESPONSE (IFNa response genes) and the KEGG_ANTIGEN_PROCESSING_AND_PRESENTATION (APM genes), which are available in the Molecular Signature Database (MSigDB) gene sets (**Supplementary Table 2,** https://www.gsea-msigdb.org/gsea/msigdb/index.jsp). Bioinformatic interrogation of these gene lists are shown in Supplementary Figures 1-3.

#### RNA-Seq Heatmaps for type I IFN response and APM genes

RNA-Seq data were quantitated to obtain raw tag counts at the gene level using either HTSeq (Anders et al, 2015) or featureCount (Liao et al, 2013). The raw tag count data was variance stabilizing transformed using VST function in DESeq2 R library (Love et al, 2014), and z-score of the transformed data was obtained to color code for heatmap. For clustered heatmaps, pheatmap R library was used with Euclidean distance and ward.D2 clustering options. Significance of gene expression changes was evaluated using DESeq2 R library and determined based on Wald-statistics (FDR<0.1) and shrunken log2 fold-change (>log2(1.3), <-log2(1.3)) using ahsr method (Stephens et al, 2017) available from DESeq2 library.

#### IPA analysis of DEGs

Ingenuity Pathway Analysis (IPA) was used to identify significantly enriched canonical pathways based on DEGs. To obtain manageable number of DEGs for IPA core analysis, different FDR and LFC were applied; siSMYD3 vs. HN6 (991 DEGs, FDR<0.05, LFC < -1 or LFC > 1), SMYD3 ASOs vs. HN6 (1401 DEGs, FDR<0.05), CRISPR KO vs. HN6 (3714 DEGs, FDR<0.01). Core analysis was performed using default options.

#### GSEA analysis of TCGA data

For gene-set enrichment analysis (GSEA), TCGA 422 HPV-negative tumors samples (Firehose Legacy) with mRNA expression data (data_RNA_Seq_v2_mRNA_median_Zscores.txt) were analyzed. A ranked gene list based on the Pearson’s correlation of each gene with SMYD3 expression was used as the ranked input to GSEA against MsigDB Hallmark gene set, h.all.v7.4.symbols.gmt.

#### Expression of retrotransposons in mRNA-Seq data

For the evaluation of differences in the expression levels of retransposons and other repetitive elements, RNA-seq retrotransposon quantification and analysis was performed using homer and bedtools. Briefly, genomic coordinates (hg38) of LINE and other repetitive elements were obtained from the UCSC genome browser consortium and quantified using the annotatePeaks.pl script, by homer.

#### CUT&RUN analysis

Raw fastq files were trimmed with cutadapt 3.4 and aligned using bowtie2-2.4.4 against hg38. For spike-in control, the trimmed fastq files were aligned against the Escherichia coil MG1655 genome. Duplicated reads were removed using Picard tools 2.25.0, and normalization factors were derived based on the uniquely mapped fragments in the corresponding spike-in control data. The enriched regions with H4K20me3, H3K9me3 and H3K4me3 signals were identified using SEACR (Meers et al, 2019) with spike-in normalization, numerical threshold (0.001), and “relaxed” options. Then, any regions enriched with IgG signals were removed via differential analysis using DESeq to obtain a final list of regions for downstream analysis. Differential peaks were detected using DESeq2 (FDR<0.1).

#### Annotation of peaks (ChIP, CUT&RUN)

Each peak/enriched region in CUT&RUN assays was annotated with the nearby gene displaying the shortest distance between TSS and the center of each peak using the Homer annotatePeaks.pl function. Pre-identified immune-related genes were used to annotate heatmap. Genome browser track and genomic coordinate heatmaps were obtained using deepTools (Ramirez et al, 2016).

#### Volcano plots

For all volcano plots, EnhancedVolcano R library was used.

#### Single-cell RNA-sequencing analysis

Published single-cell RNA-seq data were obtained from and processed as described by Puram et al (2018). Processed expression data were downloaded from Gene Expression Omnibus (GSE103322) and subjected to log2 transformation after adding one to each value. Analysis was performed using R V.4.1.2; heatmaps were generated using pheatmap (R Core Team (2021). R: A language and environment for statistical computing. R Foundation for Statistical Computing, Vienna, Austria. URL https://www.R-project.org/.; Raivo Kolde (2019). pheatmap: Pretty Heatmaps. R package version 1.0.12. https://CRAN.R-project.org/package=pheatmap). Pearson correlation was used for correlation-based analysis.

#### Statistics

To compare IHC scores among normal, dysplastic buccal squamous epithelium and HNSCC, the Kruskal-Wallis test followed by Dunn’s post hoc test with Benjamini-Hochberg (BH) procedure for multiple comparisons was performed using stats and FSA R libraries. Pearson correlations between SMYD3 and CD8 and between SMYD3 and UHRF1 immunohistochemical staining was obtained using stats R library. Associations between clinical to pathological downstaging and pretreatment mRNA expression levels of SMYD3, UHRF1, and CD8 were evaluated using published mRNA data (dbGaP, phs001623). The TPM values of SMYD3, UHRF1, and CD8A mRNA expression were used to perform the Wilcoxon rank sum test between groups with (n=4) and without (n=16) clinical to pathological downstaging.

#### Survival analysis

We generated a combined variable (SMYD3.UHRF1) using binned SMYD3 and UHRF1 QuPath scores. Then, we performed Kaplan Meier (KM) progression-free survival (PFS) and overall survival (OS) analysis to evaluate clinical outcomes of the partitioned patient groups based on the combined variable (SMYD3.UHRF1) and CD8. To partition patient groups, we searched for the thresholds that maximize the separation of patient groups in their survival curves. For KM PFS analysis, among 31 patients with progression-free survival data, the top 80% of SMDY3 and top 65% of UHRF1 QuPath scores were designated as a “high.high” group. The rest of the patients were designated as a “low” group. CD8 QuPath scores were binned as “low” (<bottom 30%) and “high.” For KM OS analysis, among 35 patients with overall survival data, the top 20% of SMDY3 and top 25% of UHRF1 QuPath scores were designated as a “high.high” group. The rest of the patients were defined as a “low” group. CD8 QuPath scores were binned as “low” (<bottom 35%) and “high.” KM analysis was performed using survival and survminer R libraries.

## KEY RESOURCES TABLE

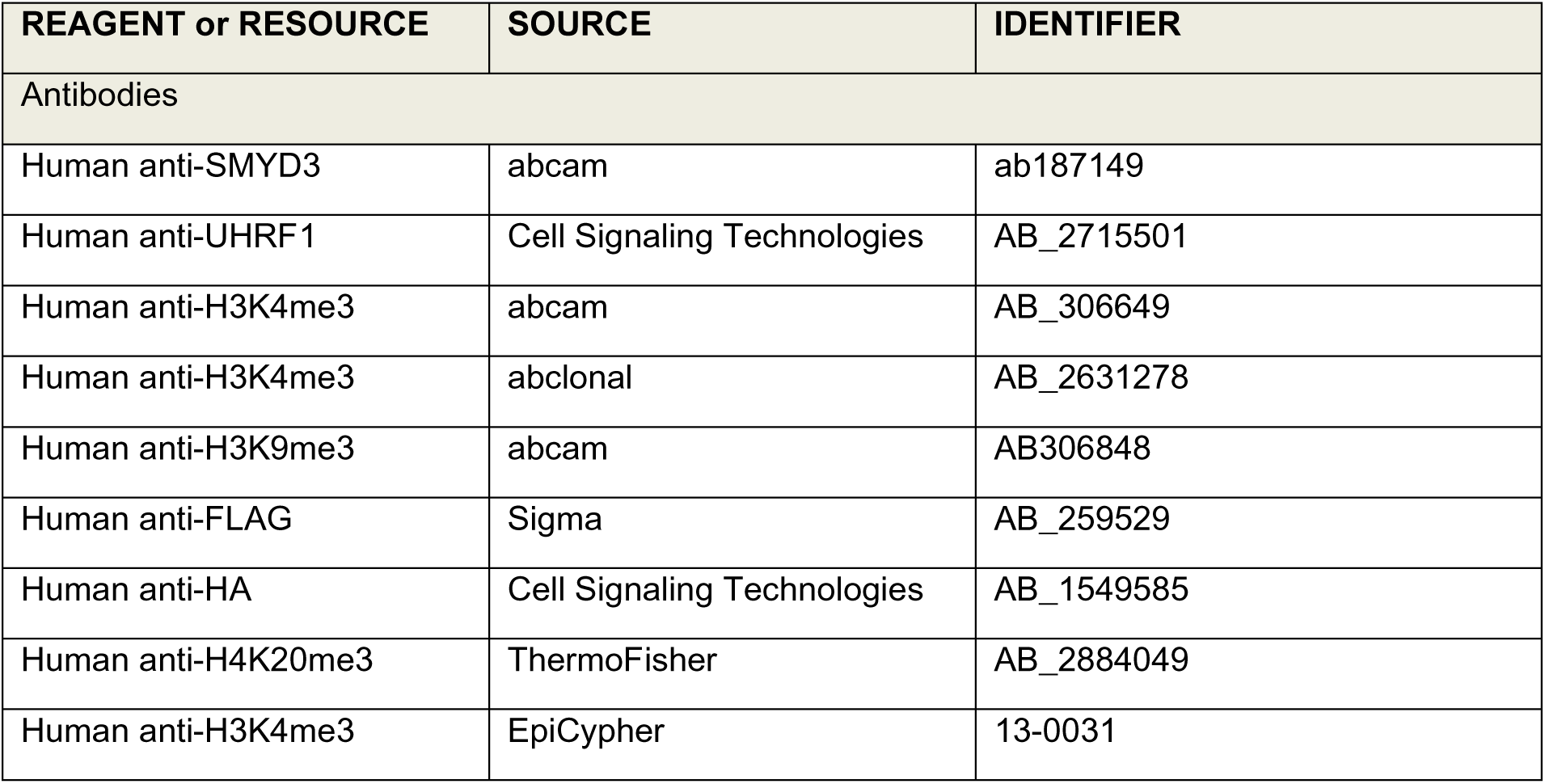
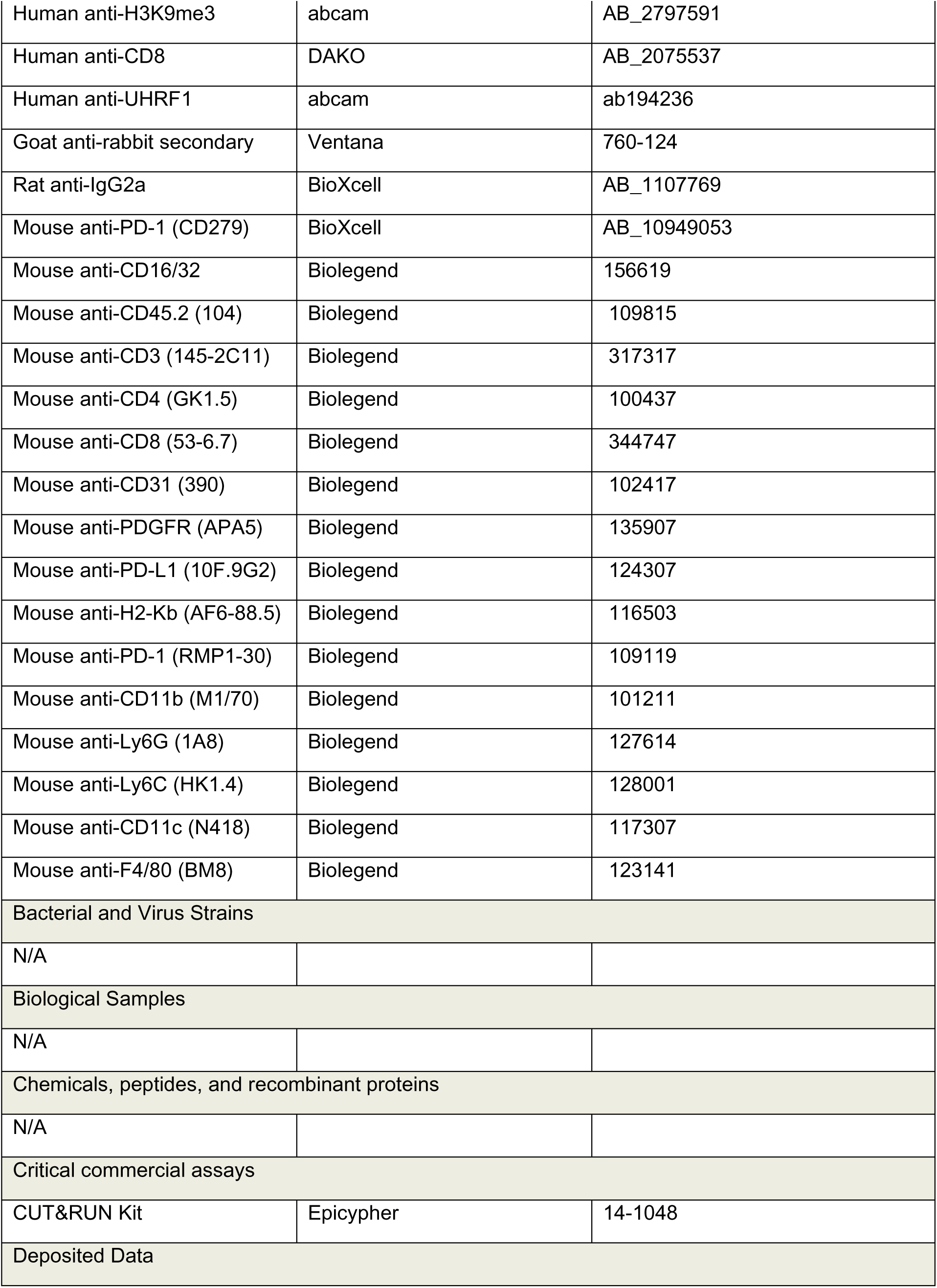
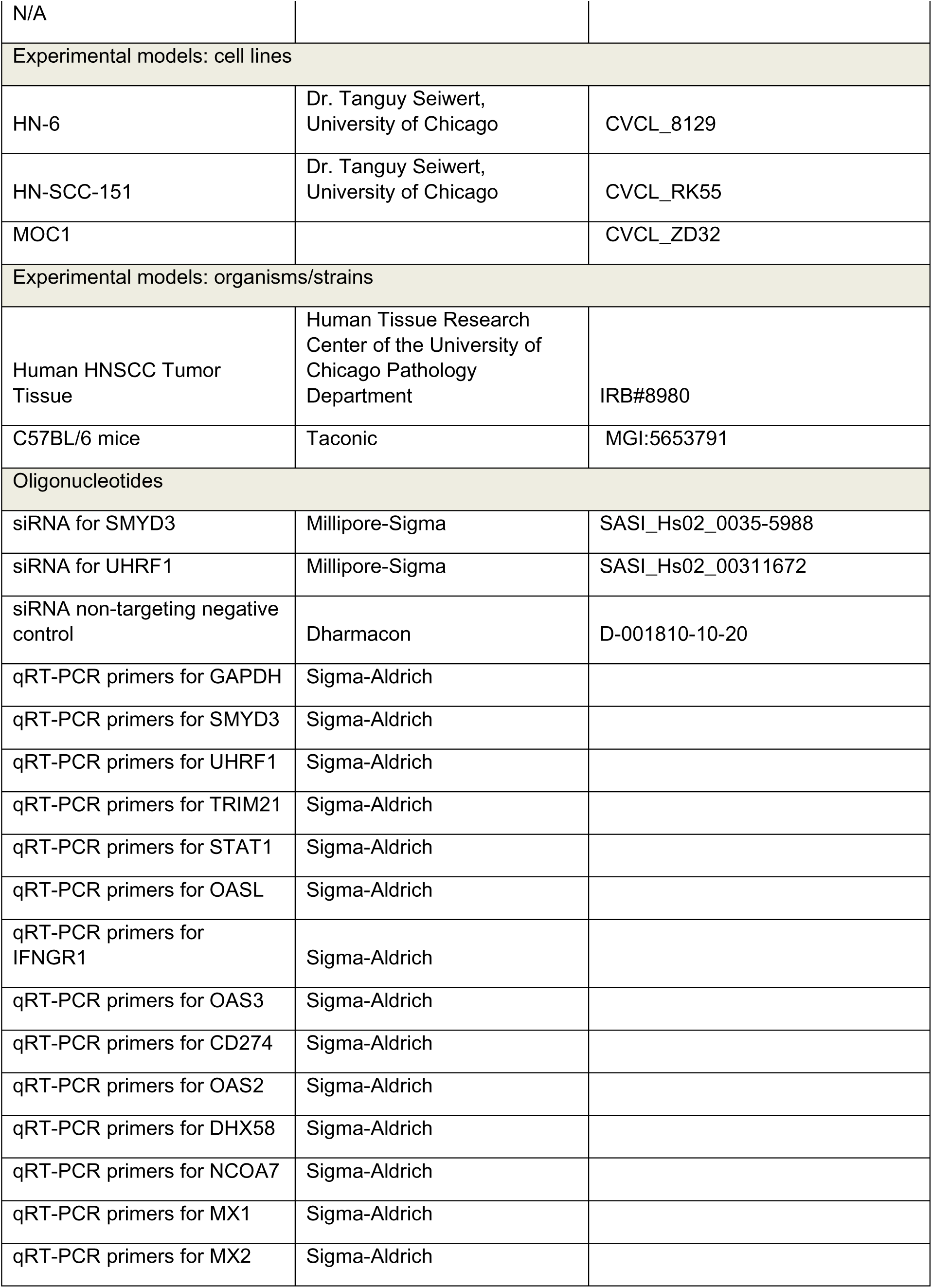
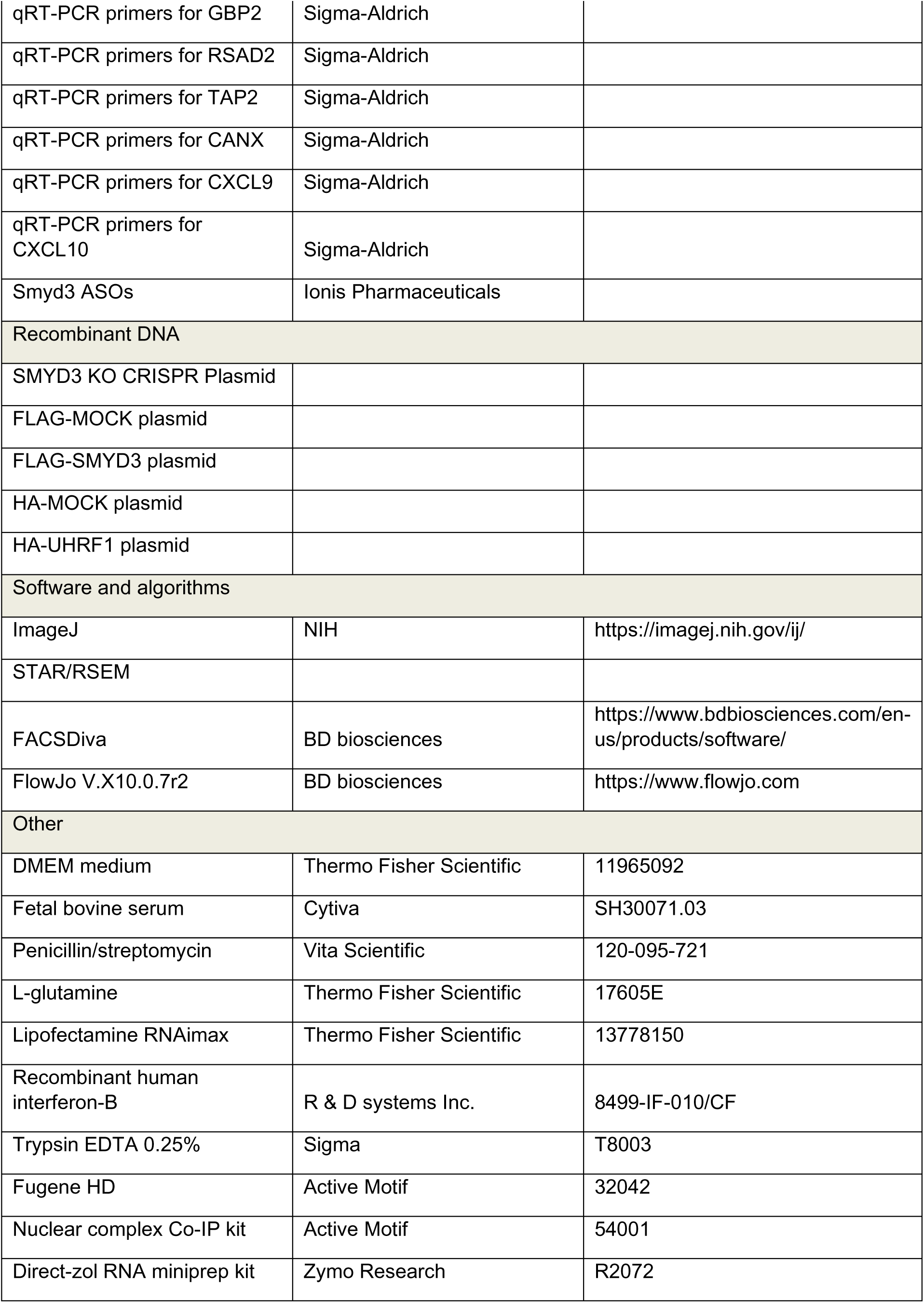
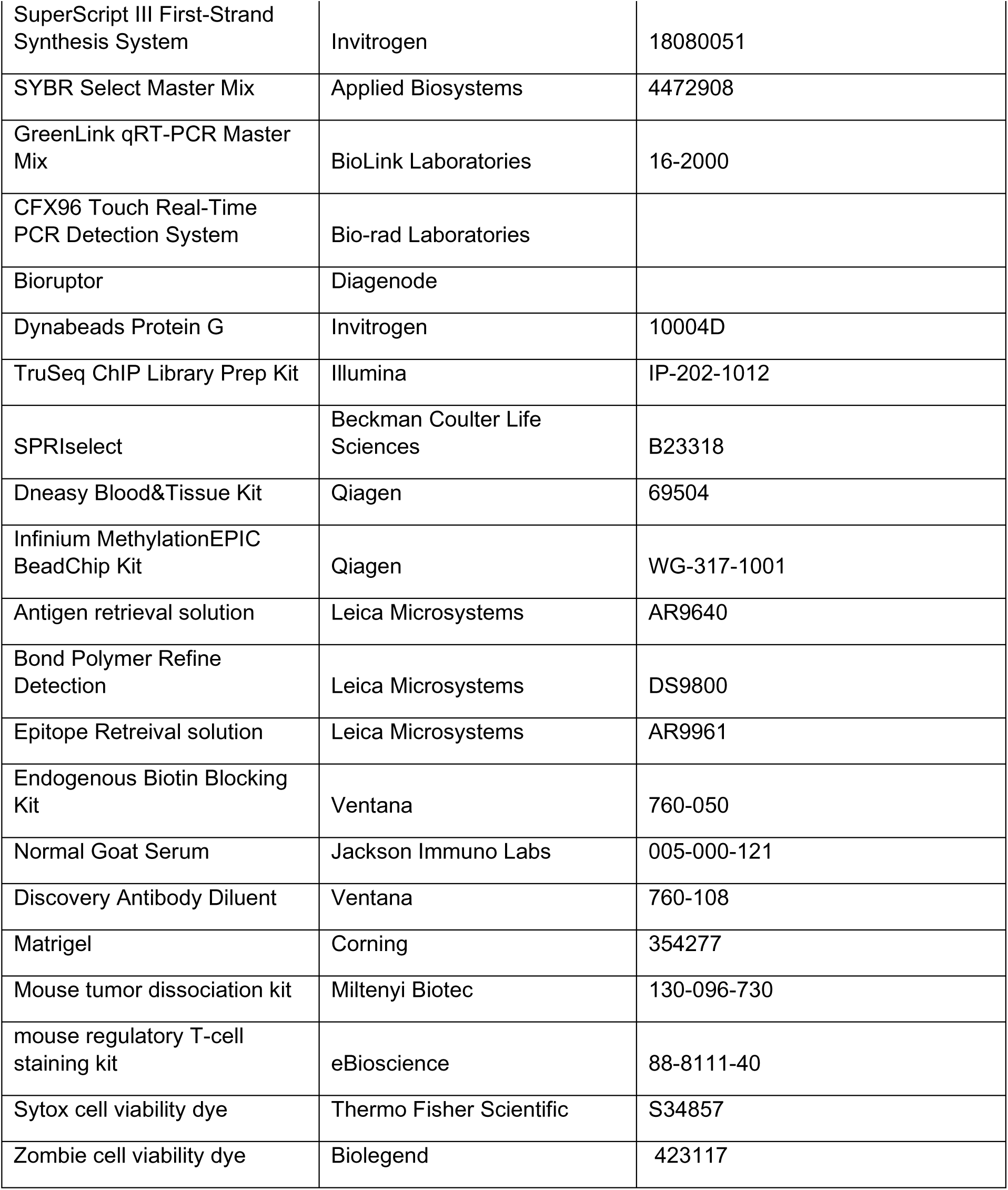

## SUPPLEMENTAL INFORMATION

**Supplementary Table 1.** List of type I IFN response, APM and MHC class I genes.

**Supplementary Table 2.** Gene set from the HALLMARK_INTERFERON_ALPHA_RESPONSE (IFNa response genes) and the KEGG_ANTIGEN_PROCESSING_AND_PRESENTATION (APM genes) (Molecular Signature Database (MSigDB)).

**Supplementary Table 3.** List of chromatin modifiers and epigenetic regulators.

**Supplementary Table 4.** Immune-related genes with significantly decreased H4K20me3 peaks in promoters/TSS/gene body regions.

**Supplementary Figure 1A, B.** Comprehensive RNA-seq heatmaps of HN-6 cells treated with siSMYD3 for 72h and IFN-β exposure for 24h; (A) IFNa GSEA gene set, (B) APM GSEA gene set.

**Supplementary Figure 2A, B.** Comprehensive RNA-seq heatmaps of HN-6 cells treated with PBS or SMYD3 ASOs for 72h and IFN-β exposure for 24h; (A) IFNa GSEA gene set, (B) APM GSEA gene set.

**Supplementary Figure 3A, B.** Comprehensive RNA-seq heatmaps of a CRISPR SMYD3 KO cell line (clone 5-3) exposed to IFN-β for 24h; (A) IFNa GSEA gene set, (B) APM GSEA gene set.

**Supplementary Figure 4.** Volcano plots showing DESeq2 results in SMYD3 ASO treated HN-6 cells **(A)** and siSMYD3 treated HN-6 cells **(B)** for 72h, exposed to IFN-β for 24h. FDR: 0.1, log2FC threshold: log2 (1.3). Red triangles: IFNa genes (from GSEA gene set), blue crosses: APM genes (from GSEA gene set), gray circles: other genes.

**Supplementary Figure 5.** Number of common, significantly (FDR<0.1, log2 fold change>abs(log2(1.3)) upregulated IFNa response genes among 97 Hallmark IFNa response genes (GSEA).

**Supplementary Figure 6.** Gene Set Enrichment Analysis (GSEA) reveals enrichment of pathways related to inflammation in an HPV-negative cell line (HN-6) after SMYD3 depletion with SMYD3 ASOs for 72h and IFN-β exposure for 24h.

**Supplementary Figure 7.** Single cell RNA seq analysis in HPV-negative HNSCC cancer cells associating SMYD3 with APM genes.

**Supplementary Figure 8.** Volcano plots of RNA-seq (left panel) and H3K4me3 ChIP-seq (right panel) in HN-6 cells after treatment with negative control or SMYD3-targeting siRNA for 3 days. Dots represent H3K4me3 peaks (35,981 variables). Log2 FC: log2 fold change>1.3. UHRF1 highlighted in purple font.

**Supplementary Figure 9. (A)** FLAG-ChIP followed by qPCR for the *UHRF1* transcription start site (TSS) or an intergenic region (IG) of *HLA-F*, designated as a negative control binding site for SMYD3. HN-6 cells were transfected for 48h with FLAG-Mock or FLAG-SMYD3 and ChIP was conducted using an anti-FLAG antibody. Enrichment as a percent of the input was calculated for each ChIP sample using the formula: 100∗ 2^[CtInput – log2(800/30)-Ct IP], where 800/30 is the input dilution factor. In this experiment, Ct values of 2 technical replicates and 2 biological replicates were used for the analysis. We observed FLAG-SMYD3 enrichment similar to background noise near the TSS of the *UHRF1* locus in all the replicates tested. **(B)** USCS tracks of Smyd3 ChIP in mouse hepatocellular carcinoma cells focusing on the mouse *Uhrf1* gene locus (Sarris et al). Tracks show enrichment of Smyd3 at the *Uhrf1* TSS.

**Supplementary Figure 10.** Upregulation of immune-related genes after UHRF1 depletion in the absence of IFN-β. HN-SCC-151 cells were transfected with control versus UHRF1-targeting siRNAs for 72h. Cells were collected for RNA extraction and cDNA synthesis. SYBR green qPCR was conducted. Similar results were obtained with two biological replicates.

**Supplementary Figure 11.** Western blotting for SMYD3 and UHRF1 in HN-6 and 5-2 SMYD3 KO cells. Cells were collected and nuclear extraction was conducted followed by Western blotting for SMYD3 and UHRF1. 10ug for SMYD3 and 15ug for UHRF1 of nuclear extract were loaded for all conditions. H3 was used as a loading control.

**Supplementary Figure 12. Top heatmap:** RNA expression z-score heatmap of 26 immune-related genes corresponding to 38 differential H4K20me3 peaks present on respective promoters, TSS or gene body regions in HN-6 cells and CRISPR SMYD3 KO 5-3 cells after 24h of exposure to IFN-β (P adj <0.1). **Bottom heatmap:** Expression z-score of CUT&RUN signals corresponding to H4K20me3 peaks present in promoters, TSS or gene body regions in SMYD3 KO (5-3 cells) compared to HN-6. Peaks in the heatmap are ordered the same as in the RNA expression z-score heatmap.

**Supplementary Figure 13. CUT&RUN assay for H3K4me3 and H3K9me3 in SMYD3 KO 5-3 cells compared to control HN-6 cells after 24h exposure to IFN-β.** Volcano plot of H3K4me3 peaks **(A)** and heatmap of H3K4me3 peaks corresponding to a panel of representative type I IFN response genes **(B)**. Volcano plot of H3K9me3 peaks **(C)** and heatmap of H3K9me3 peaks corresponding to a panel of representative type I IFN response genes **(D)**.

**Supplementary Figure 14. RNA-seq retrotransposon quantification and analysis was performed using homer and bedtools.** HN-6 cells were treated with siNC or siSMYD3 for 3 days and for 24h with IFN-β prior to collection. RNA was extracted and mRNA-seq was performed. RPKM; per million mapped reads.

**Supplementary Figure 15.** Western blotting for Smyd3 in MOC1 cells treated with increasing molar concentrations of control or Smyd3 targeting ASOs. MOC1 cells were plated in 10cm dishes and treated with 0, 0.5, 0.1, 0.02 or 0.004uM of control ASOs or Smyd3 ASOs for 72h. Cells were collected and nuclear extraction was conducted. 10ug of nuclear extract were loaded for Smyd3 blotting, and H3 was used as a loading control. 0.5uM of Smyd3 ASOs induced near complete knockdown of Smyd3.

**Supplementary Figure 16. (A)** Comparison of SMYD3 protein expression levels in the cancer cell versus stroma cell compartment. H-score was determined by QuPath. **(B)** Mean SMYD3 mRNA expression levels in different cell types assessed from a publicly available single-cell RNA-seq database. Wilcoxon rank sum test with continuity correction, p<0.0001.

**Supplementary Figure 17.** Correlation between UHRF1 and CD8 protein levels in 64 HPV-negative HNSCC tumors. Pearson’s correlation coefficient rho *R*=-0.011, p=093.

**Supplementary Figure 18.** SMYD3 (A), UHRF1 (B) QuPath Score and CD8 % positivity (C) and Kaplan Meier curves for overall survival.

**Supplementary Figure 19.** mRNA expression levels of STING1 after SMYD3 depletion using siRNAs **(A)** or CRISPR (5-3 cell line) **(B)** and 24h of IFN-β exposure.

**Supplementary Figure 20.** Genomic alterations of SMYD3 in HPV-negative tumors samples of the TCGA dataset (Firehose Legacy).

**Supplementary Figure 21.** Western blotting of CRISPR SMYD3 KO cell lines 5-2 and 5-3. 10ug of nuclear extracts were loaded and blotted for SMYD3. H3 was used as a loading control.

