## Supplementary Figures for "Epigenetic silencing by SMYD3 represses tumor intrinsic interferon response in HPV-negative squamous cell carcinoma of the head and neck"

**Supplementary Figure 1A, B.** Comprehensive RNA-seq heatmaps of HN-6 cells treated with siSMYD3 for 72h and IFN-β exposure for 24h; (A) IFNa GSEA gene set, (B) APM GSEA gene set.

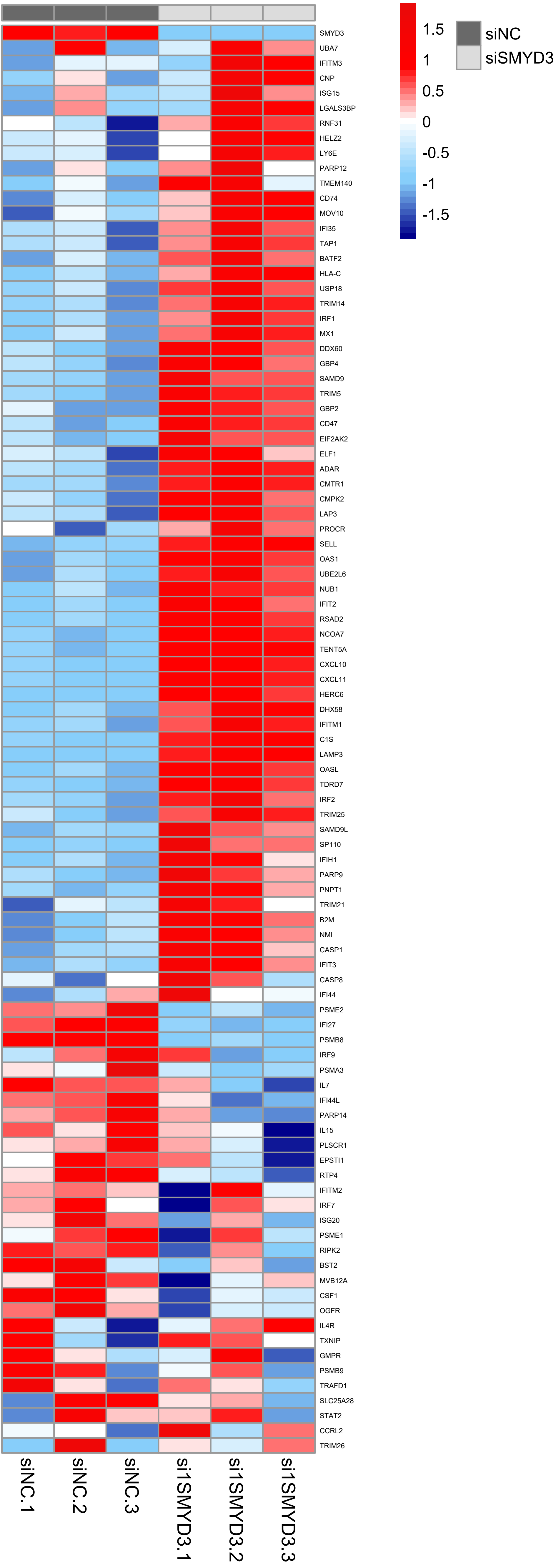
1.5
1
0.5
0
-0.5
-1
-1.5
siNC
siSMYD3
SMYD3
UBA7
IFITM3
CNP
ISG15
LGALS3BP
RNF31
HELZ2
LY6E
PARP12
TMEM140
CD74
MOV10
IFI35
TAP1
BATF2
HLA-C
USP18
TRIM14
IRF1
MX1
DDX60
GBP4
SAMD9
TRIM5
GBP2
CD47
EIF2AK2
ELF1
ADAR
CMTR1
CMPK2
LAP3
PROCR
SELL
OAS1
UBE2L6
NUB1
IFIT2
RSAD2
NCOA7
TENT5A
CXCL10
CXCL11
HERC6
DHX58
IFITM1
C1S
LAMP3
OASL
TDRD7
IRF2
TRIM25
SAMD9L
SP110
IFIH1
PARP9
PNPT1
TRIM21
B2M
NMI
CASP1
IFIT3
CASP8
IFI44
PSME2
IFI27
PSMB8
IRF9
PSMA3
IL7
IFI44L
PARP14
IL15
PLSCR1
EPSTI1
RTP4
IFITM2
IRF7
ISG20
PSME1
RIPK2
BST2
MVB12A
CSF1
OGFR
IL4R
TXNIP
GMPR
PSMB9
TRAFD1
SLC25A28
STAT2
CCRL2
TRIM26
siNC.1
siNC.2
siNC.3
si1SMYD3.1
si1SMYD3.2
si1SMYD3.3

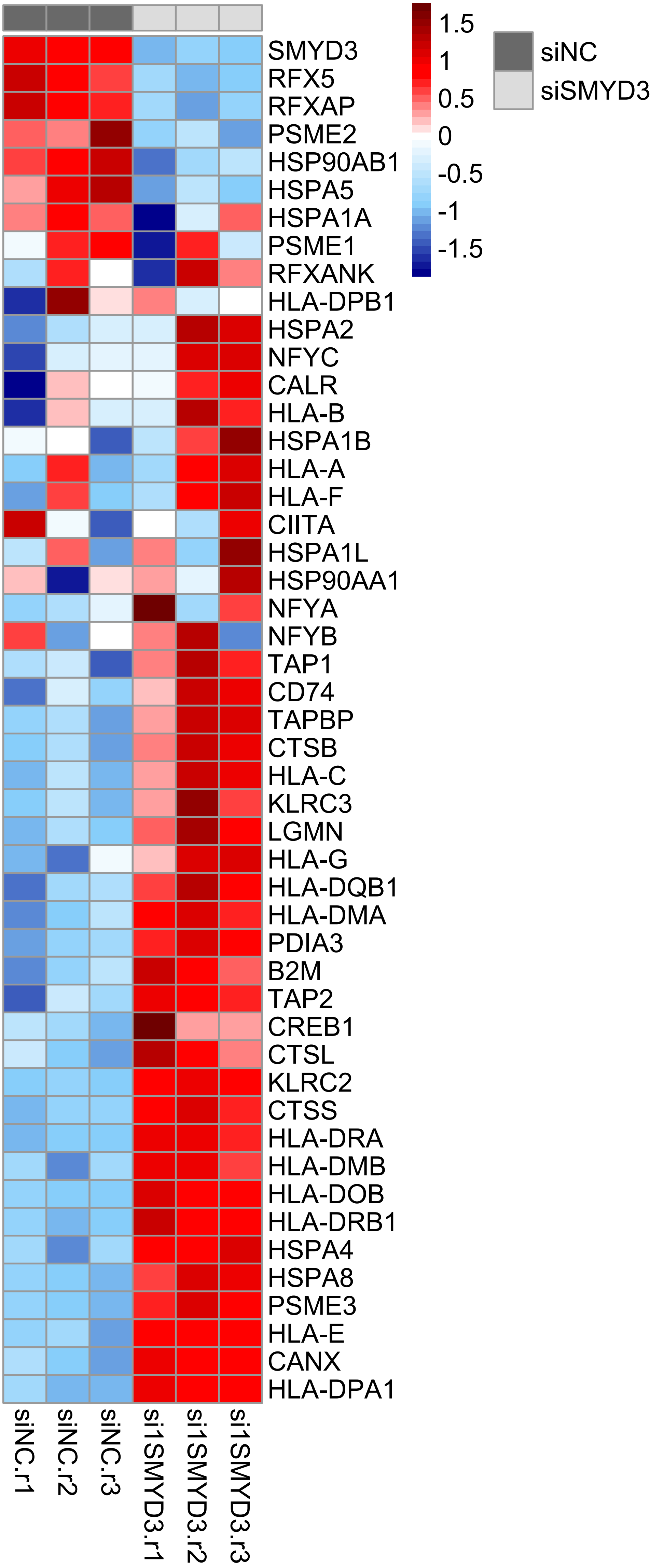
siNC
siSMYD3
1.5
1
0.5
0
-0.5
-1
-1.5
SMYD3
RFX5
RFXAP
PSME2
HSP90AB1
HSPA5
HSPA1A
PSME1
RFXANK
HLA-DPB1
HSPA2
NFYC
CALR
HLA-B
HSPA1B
HLA-A
HLA-F
CIITA
HSPA1L
HSP90AA1
NFYA
NFYB
TAP1
CD74
TAPBP
CTSB
HLA-C
KLRC3
LGMN
HLA-G
HLA-DQB1
HLA-DMA
PDIA3
B2M
TAP2
CREB1
CTSL
KLRC2
CTSS
HLA-DRA
HLA-DMB
HLA-DOB
HLA-DRB1
HSPA4
HSPA8
PSME3
HLA-E
CANX
HLA-DPA1
siNC.r1
siNC.r2
siNC.r3
si1SMYD3.r1
si1SMYD3.r2
si1SMYD3.r3

**Supplementary Figure 2A, B.** Comprehensive RNA-seq heatmaps of HN-6 cells treated with PBS or SMYD3 ASOs for 72h and IFN-β exposure for 24h; (A) IFNa GSEA gene set, (B) APM GSEA gene set.

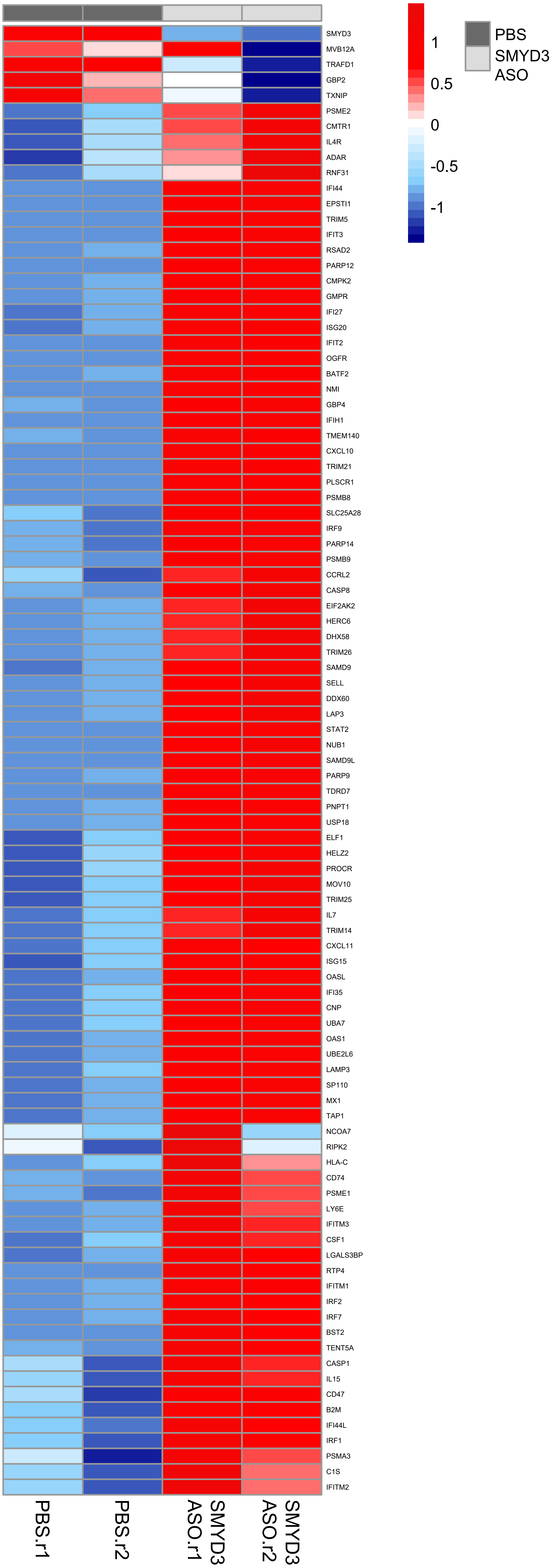
PBS
SMYD3 ASO
1
0.5
0
-0.5
-1
SMYD3
MVB12A
TRAFD1
GBP2
TXNIP
PSME2
CMTR1
IL4R
ADAR
RNF31
IFI44
EPSTI1
TRIM5
IFIT3
RSAD2
PARP12
CMPK2
GMPR
IFI27
ISG20
IFIT2
OGFR
BATF2
NMI
GBP4
IFIH1
TMEM140
CXCL10
TRIM21
PLSCR1
PSMB8
SLC25A28
IRF9
PARP14
PSMB9
CCRL2
CASP8
EIF2AK2
HERC6
DHX58
TRIM26
SAMD9
SELL
DDX60
LAP3
STAT2
NUB1
SAMD9L
PARP9
TDRD7
PNPT1
USP18
ELF1
HELZ2
PROCR
MOV10
TRIM25
IL7
TRIM14
CXCL11
ISG15
OASL
IFI35
CNP
UBA7
OAS1
UBE2L6
LAMP3
SP110
MX1
TAP1
NCOA7
RIPK2
HLA-C
CD74
PSME1
LY6E
IFITM3
CSF1
LGALS3BP
RTP4
IFITM1
IRF2
IRF7
BST2
TENT5A
CASP1
IL15
CD47
B2M
IFI44L
IRF1
PSMA3
C1S
IFITM2
PBS.r1
PBS.r2
SMYD3 ASO.r1
SMYD3 ASO.r2

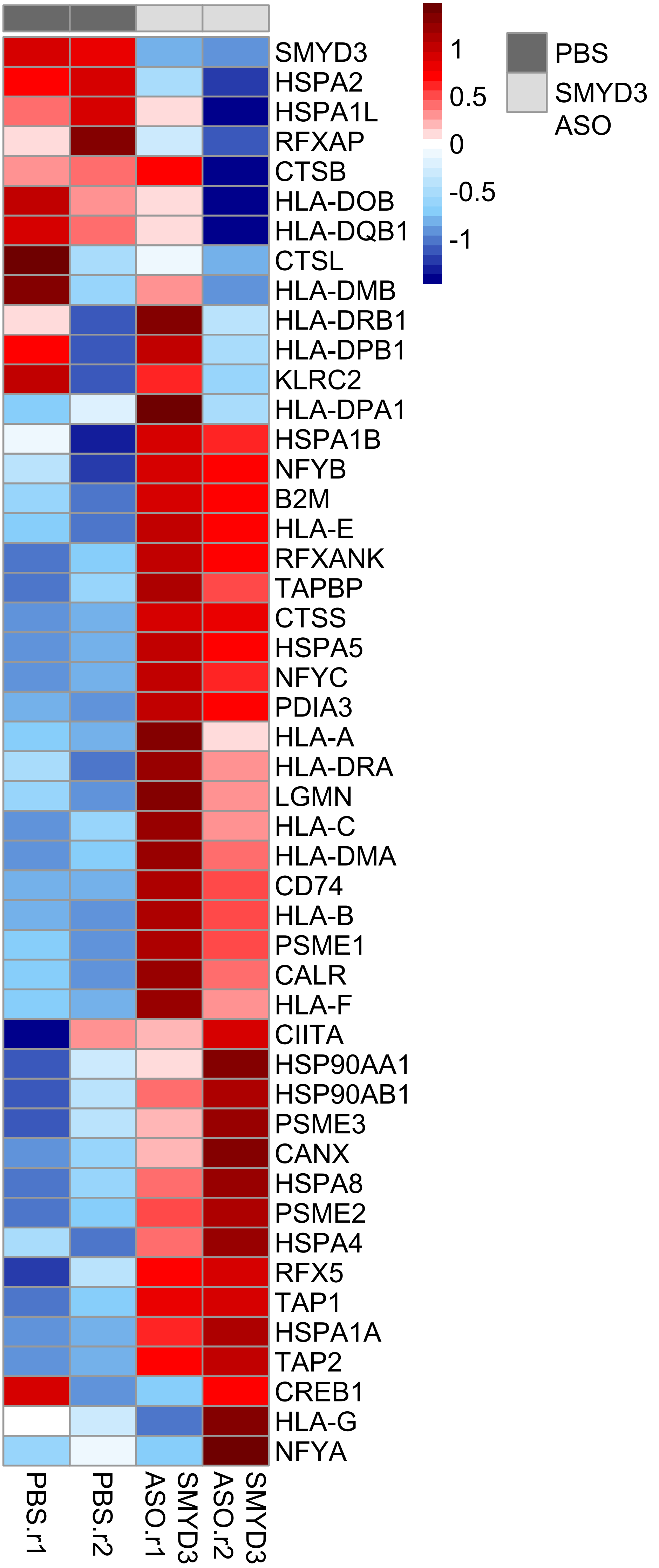

SMYD3
HSPA2
HSPA1L
RFXAP
CTSB
HLA-DOB
HLA-DQB1
CTSL
HLA-DMB
HLA-DRB1
HLA-DPB1
KLRC2
HLA-DPA1
HSPA1B
NFYB
B2M
HLA-E
RFXANK
TAPBP
CTSS
HSPA5
NFYC
PDIA3
HLA-A
HLA-DRA
LGMN
HLA-C
HLA-DMA
CD74
HLA-B
PSME1
CALR
HLA-F
CIITA
HSP90AA1
HSP90AB1
PSME3
CANX
HSPA8
PSME2
HSPA4
RFX5
TAP1
HSPA1A
TAP2
CREB1
HLA-G
NFYA
PBS.r1
PBS.r2
SMYD3 ASO.r1
SMYD3 ASO.r2
1
0.5
0
-0.5
-1
PBS
SMYD3 ASO

**Supplementary Figure 3A, B.** Comprehensive RNA-seq heatmaps of a CRISPR SMYD3 KO cell line (clone 5-3) exposed to IFN-β for 24h; (A) IFNa GSEA gene set, (B) APM GSEA gene set.

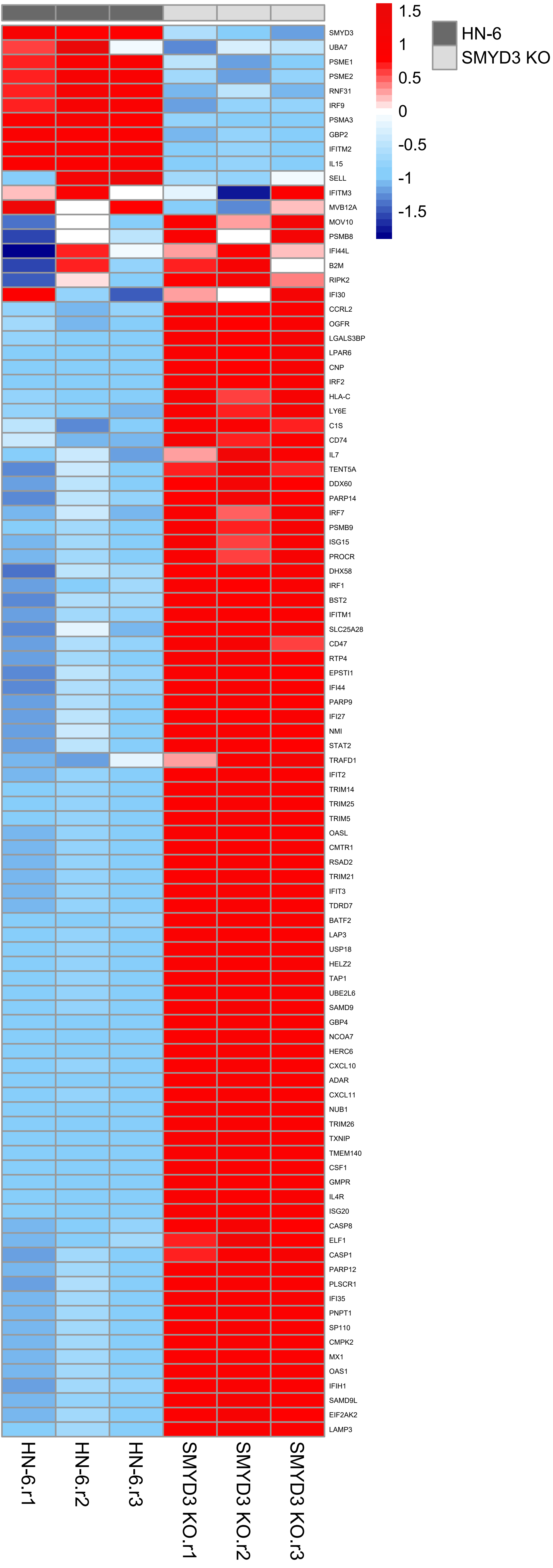

1.5
1
0.5
0
-0.5
-1
-1.5
HN-6
SMYD3 KO
SMYD3
UBA7
PSME1
PSME2
RNF31
IRF9
PSMA3
GBP2
IFITM2
IL15
SELL
IFITM3
MVB12A
MOV10
PSMB8
IFI44L
B2M
RIPK2
IFI30
CCRL2
OGFR
LGALS3BP
LPAR6
CNP
IRF2
HLA-C
LY6E
C1S
CD74
IL7
TENT5A
DDX60
PARP14
IRF7
PSMB9
ISG15
PROCR
DHX58
IRF1
BST2
IFITM1
SLC25A28
CD47
RTP4
EPSTI1
IFI44
PARP9
IFI27
NMI
STAT2
TRAFD1
IFIT2
TRIM14
TRIM25
TRIM5
OASL
CMTR1
RSAD2
TRIM21
IFIT3
TDRD7
BATF2
LAP3
USP18
HELZ2
TAP1
UBE2L6
SAMD9
GBP4
NCOA7
HERC6
CXCL10
ADAR
CXCL11
NUB1
TRIM26
TXNIP
TMEM140
CSF1
GMPR
IL4R
ISG20
CASP8
ELF1
CASP1
PARP12
PLSCR1
IFI35
PNPT1
SP110
CMPK2
MX1
OAS1
IFIH1
SAMD9L
EIF2AK2
LAMP3
HN-6.r1
HN-6.r2
HN-6.r3
SMYD3 KO.r1
SMYD3 KO.r2
SMYD3 KO.r3

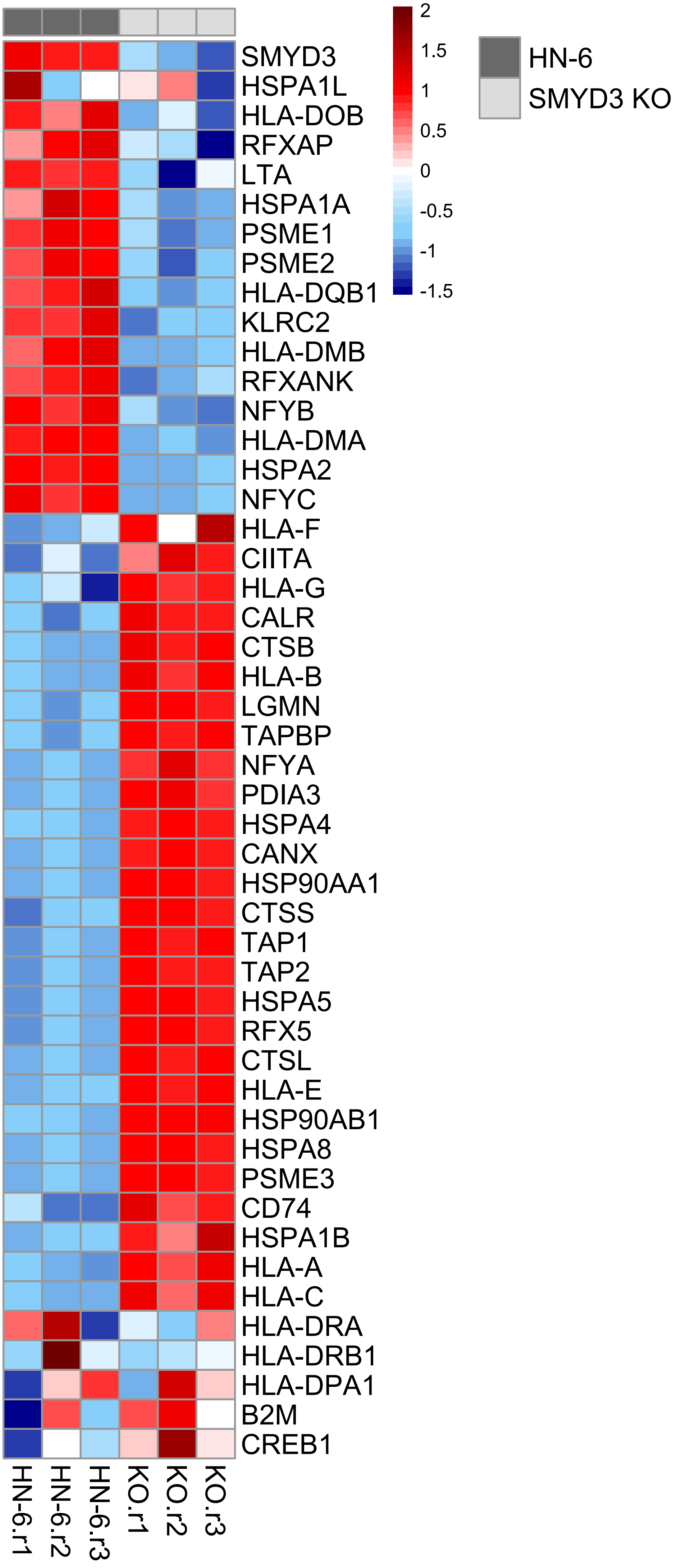

SMYD3
HSPA1L
HLA-DOB
RFXAP
LTA
HSPA1A
PSME1
PSME2
HLA-DQB1
KLRC2
HLA-DMB
RFXANK
NFYB
HLA-DMA
HSPA2
NFYC
HLA-F
CIITA
HLA-G
CALR
CTSB
HLA-B
LGMN
TAPBP
NFYA
PDIA3
HSPA4
CANX
HSP90AA1
CTSS
TAP1
TAP2
HSPA5
RFX5
CTSL
HLA-E
HSP90AB1
HSPA8
PSME3
CD74
HSPA1B
HLA-A
HLA-C
HLA-DRA
HLA-DRB1
HLA-DPA1
B2M
CREB1
HN-6.r1
HN-6.r2
HN-6.r3
KO.r1
KO.r2
KO.r3
2
1.5
1
0.5
0
-0.5
-1
-1.5
HN-6
SMYD3 KO

**Supplementary Figures 4-7**

**Supplementary Figure 4.** Volcano plots showing DESeq2 results in SMYD3 ASO treated HN-6 cells **(A)** and siSMYD3 treated HN-6 cells **(B)** for 72h, exposed to IFN-β for 24h. FDR: 0.1, log2FC threshold: log2 (1.3). Red triangles: IFNa genes (from GSEA gene set), blue crosses: APM genes (from GSEA gene set), gray circles: other genes.

**(A) SMYD3 ASO volcano plot:**

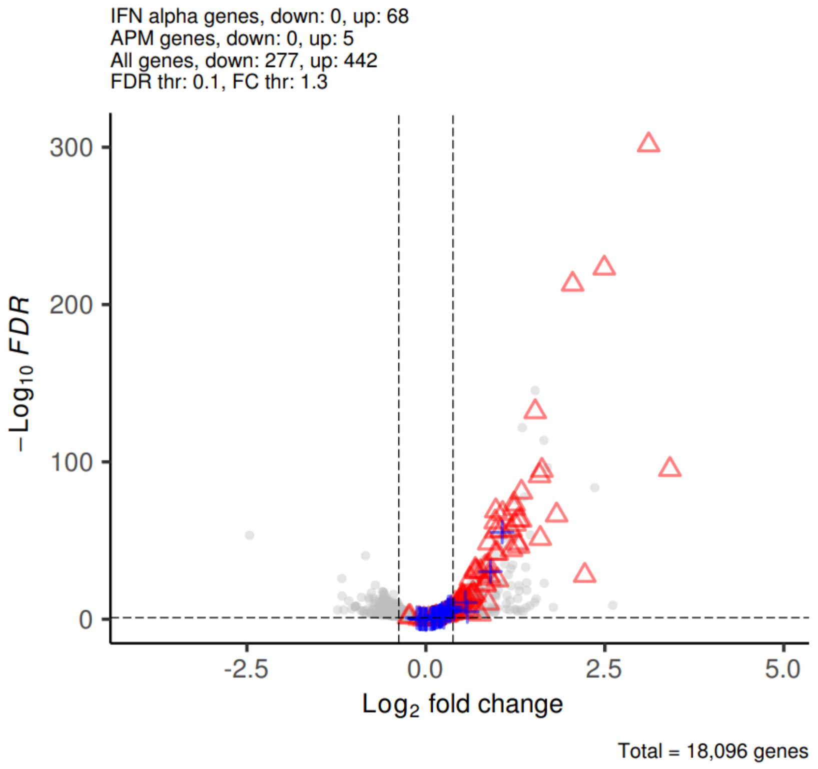


**(B) siSMYD3 volcano plot:**

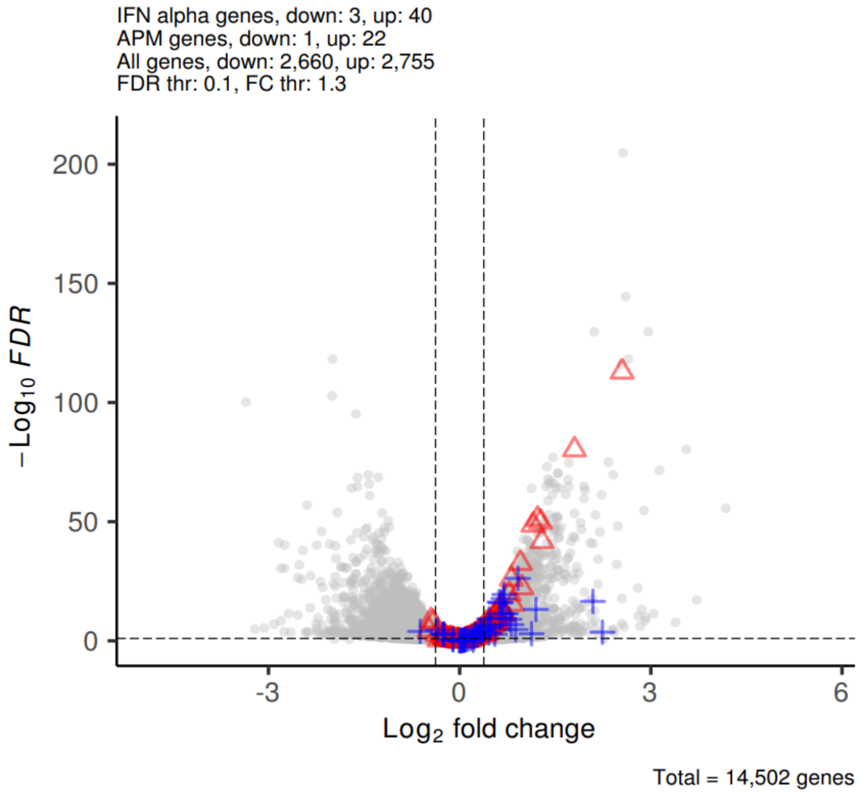


**Supplementary Figure 5.** Number of common, significantly (FDR<0.1, log2 fold change>abs(log2(1.3)) upregulated IFNa response genes among 97 Hallmark IFNa response genes (GSEA).

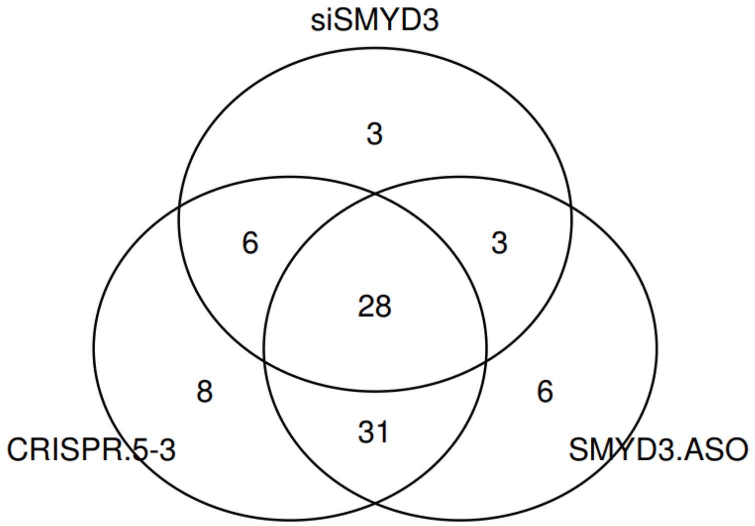

**Supplementary Figure 6.** Gene Set Enrichment Analysis (GSEA) reveals enrichment of pathways related to inflammation in an HPV-negative cell line (HN-6) after SMYD3 depletion with SMYD3 ASOs **(A)** and siSMYD3 **(B)** for 72h and IFN-β exposure for 24h.

**(A)**

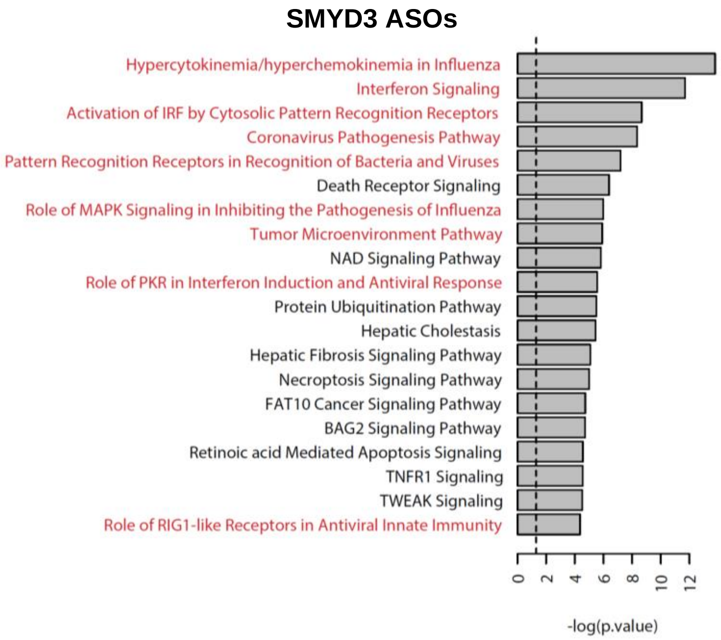


**(B)**

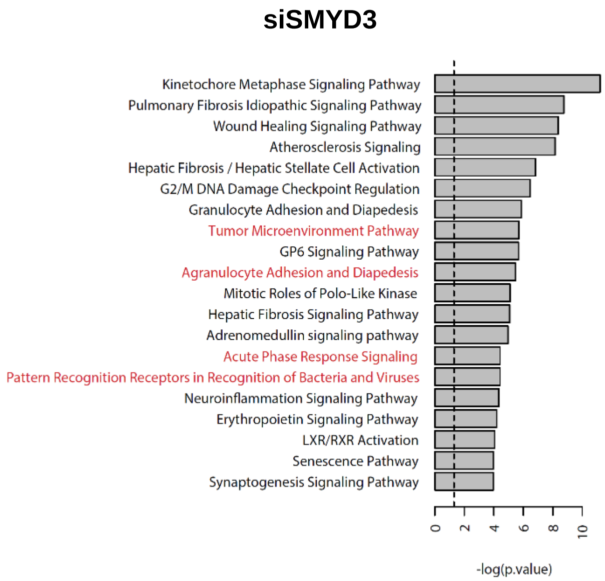

**Supplementary Figure 7.** Single cell RNA seq analysis in HPV-negative HNSCC cancer cells associating SMYD3 with APM genes.

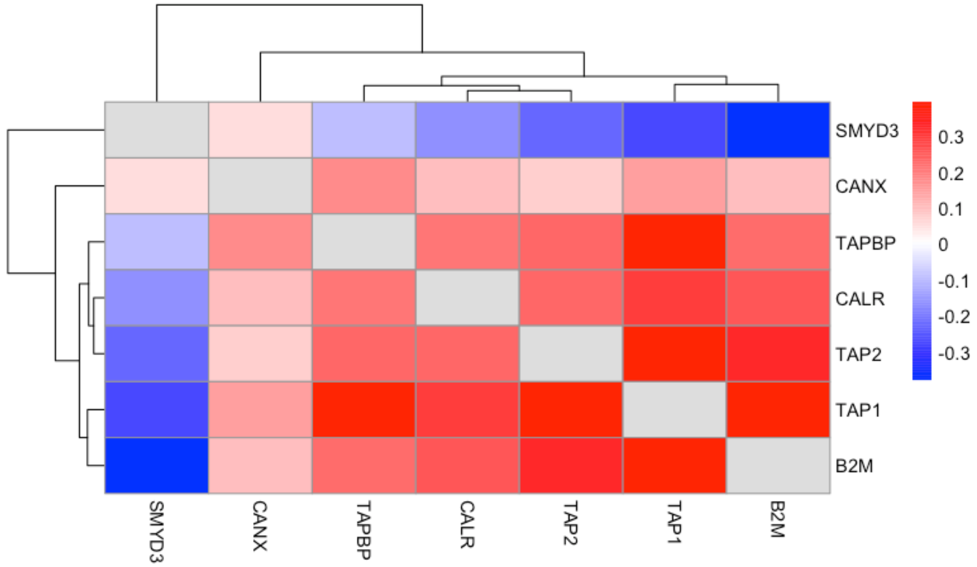

**Supplementary Figures 8-21**

**Supplementary Figure 8.** Volcano plot of RNA-seq in HN-6 cells after treatment with negative control or SMYD3-targeting siRNA for 3 days. Dots represent H3K4me3 peaks (35,981 variables). Log2 FC: log2 fold change>1.3. UHRF1 highlighted in purple font.

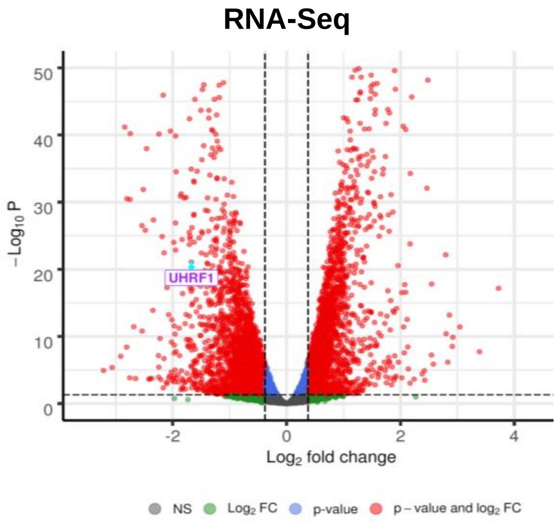


**Supplementary Figure 9.** FLAG-ChIP followed by qPCR for the *UHRF1* transcription start site (TSS) or an intergenic region (IG) of *HLA-F*, designated as a negative control binding site for SMYD3. HN-6 cells were transfected for 48h with FLAG-Mock or FLAG-SMYD3 and ChIP was conducted using an anti-FLAG antibody. Enrichment as a percent of the input was calculated for each ChIP sample using the formula: 100∗ 2ˆ[CtInput – log2(800/30)-Ct IP], where 800/30 is the input dilution factor. In this experiment, Ct values of 2 technical replicates and 2 biological replicates were used for the analysis. We observed FLAG-SMYD3 enrichment similar to background noise near the TSS of the *UHRF1* locus in all the replicates tested.

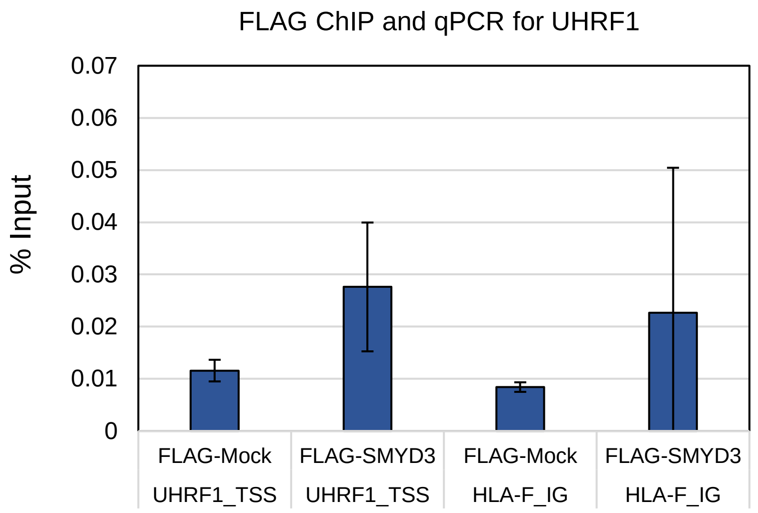


**Supplementary Figure 10.** Upregulation of immune-related genes after UHRF1 depletion in the absence of IFN-β. HN-SCC-151 cells were transfected with control versus UHRF1-targeting siRNAs for 72h. Cells were collected for RNA extraction and cDNA synthesis. SYBR green qPCR was conducted. Similar results were obtained with two biological replicates.

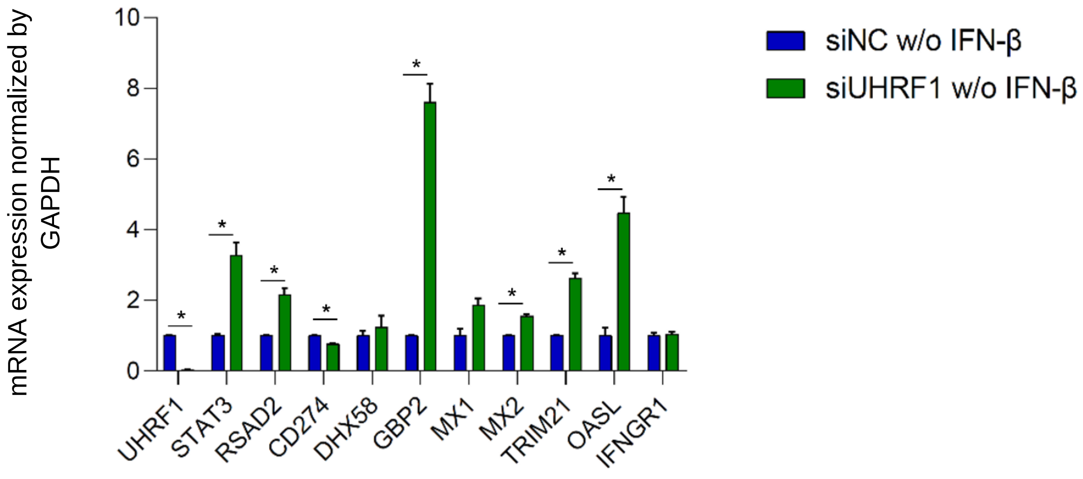

**Supplementary Figure 11.** Western blotting for SMYD3 and UHRF1 in HN-6 and 5-2 SMYD3 KO cells. Cells were collected and nuclear extraction was conducted followed by Western blotting for SMYD3 and UHRF1. 10ug for SMYD3 and 15ug for UHRF1 of nuclear extract were loaded for all conditions. H3 was used as a loading control.

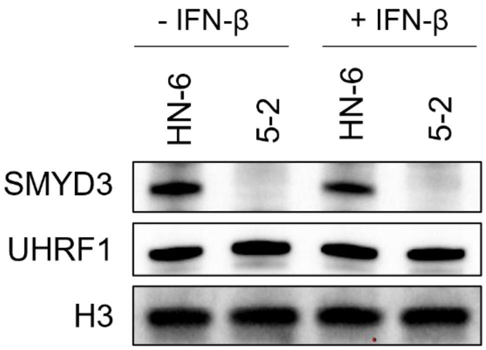


**Supplementary Figure 12. Top heatmap:** RNA expression z-score heatmap of 26 immune-related genes corresponding to 38 differential H4K20me3 peaks present on respective promoters, TSS or gene body regions in HN-6 cells and CRISPR SMYD3 KO 5-3 cells after 24h of exposure to IFN-β (P adj <0.1). **Bottom heatmap:** Expression z-score of CUT&RUN signals corresponding to H4K20me3 peaks present in promoters, TSS or gene body regions in SMYD3 KO 5-3 compared to HN-6 cells. Peaks in the heatmap are ordered the same as in the RNA expression z-score heatmap.

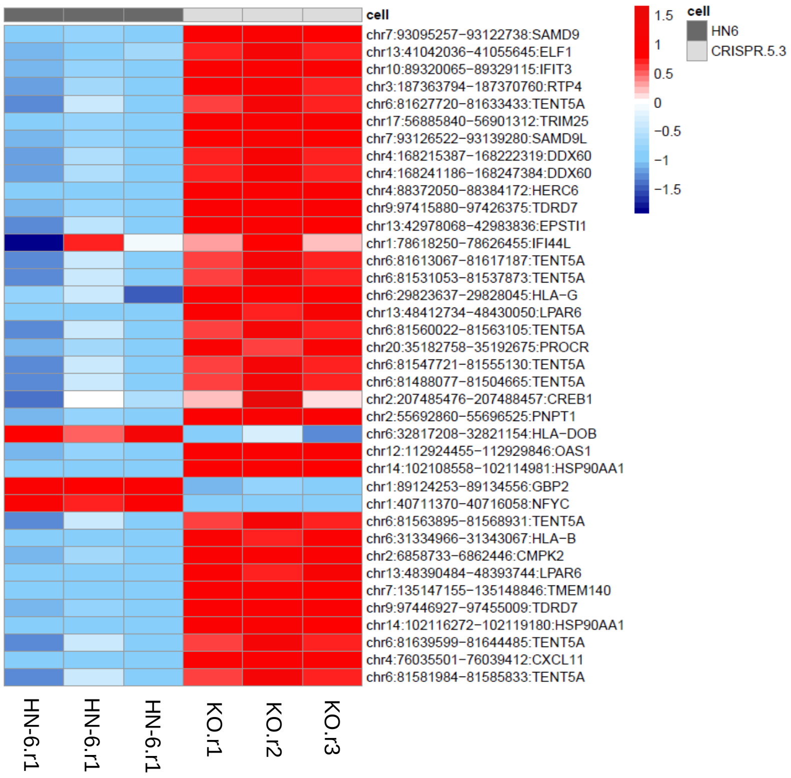

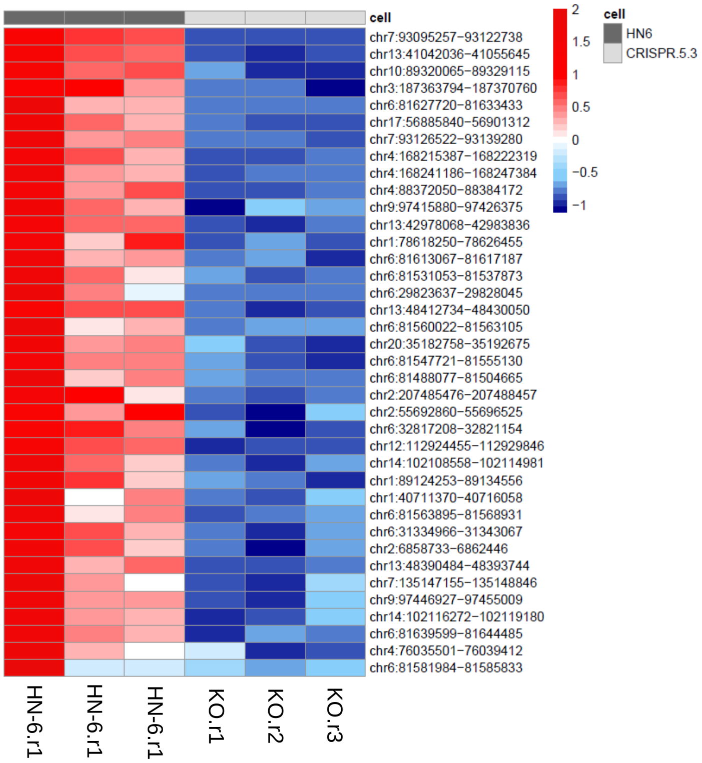
cell
chr7:93095257−93122738
chr13:41042036−41055645
chr10:89320065−89329115
chr3:187363794−187370760
chr6:81627720−81633433
chr17:56885840−56901312
chr7:93126522−93139280
chr4:168215387−168222319
chr4:168241186−168247384
chr4:88372050−88384172
chr9:97415880−97426375
chr13:42978068−42983836
chr1:78618250−78626455
chr6:81613067−81617187
chr6:81531053−81537873
chr6:29823637−29828045
chr13:48412734−48430050
chr6:81560022−81563105
chr20:35182758−35192675
chr6:81547721−81555130
chr6:81488077−81504665
chr2:207485476−207488457
chr2:55692860−55696525
chr6:32817208−32821154
chr12:112924455−112929846
chr14:102108558−102114981
chr1:89124253−89134556
chr1:40711370−40716058
chr6:81563895−81568931
chr6:31334966−31343067
chr2:6858733−6862446
chr13:48390484−48393744
chr7:135147155−135148846
chr9:97446927−97455009
chr14:102116272−102119180
chr6:81639599−81644485
chr4:76035501−76039412
chr6:81581984−81585833
HN-6.r1
HN-6.r1
HN-6.r1
KO.r1
KO.r2
KO.r3
2
1.5
1
0.5
0
−0.5
−1
cell
HN6
CRISPR.5.3

**Supplementary Figure 13. CUT&RUN assay for H3K4me3 and H3K9me3 in SMYD3 KO 5-3 cells compared to control HN-6 cells after 24h exposure to IFN-β.** Volcano plot of H3K4me3 peaks **(A)** and heatmap of H3K4me3 peaks corresponding to a panel of representative type I IFN response genes **(B)**. Volcano plot of H3K9me3 peaks **(C)** and heatmap of H3K9me3 peaks corresponding to a panel of representative type I IFN response genes **(D)**.

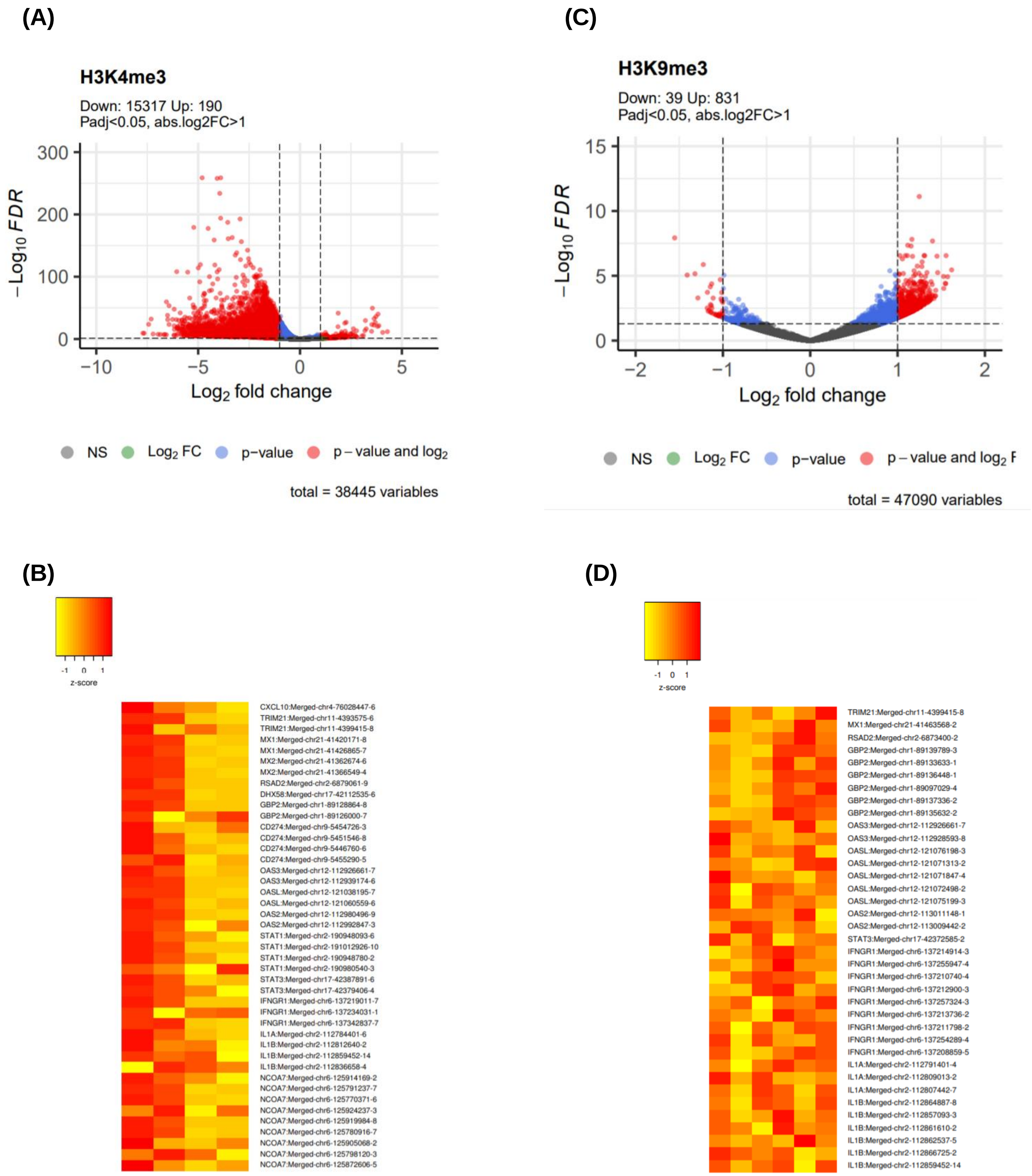

**Supplementary Figure 14. RNA-seq retrotransposon quantification and analysis was performed using homer and bedtools.** HN-6 cells were treated with siNC or siSMYD3 for 3 days and for 24h with IFN-β prior to collection. RNA was extracted and mRNA-seq was performed. RPKM; per million mapped reads.

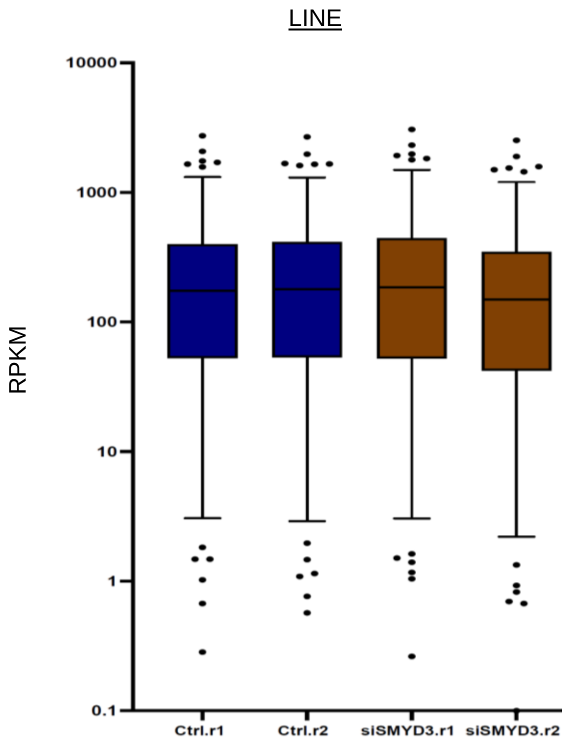

**Supplementary Figure 15.** Western blotting for Smyd3 in MOC1 cells treated with increasing molar concentrations of control or Smyd3 targeting ASOs. MOC1 cells were plated in 10cm dishes and treated with 0, 0.5, 0.1, 0.02 or 0.004uM of control ASOs or Smyd3 ASOs for 72h. Cells were collected and nuclear extraction was conducted. 10ug of nuclear extract were loaded for Smyd3 blotting, and H3 was used as a loading control. 0.5uM of Smyd3 ASOs induced near complete knockdown of Smyd3.

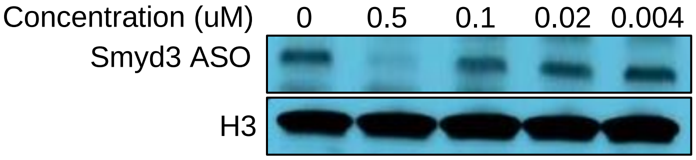


**Supplementary Figure 16. (A)** Comparison of SMYD3 protein expression levels in the cancer cell versus stroma cell compartment. H-score was determined by QuPath. Wilcoxon rank sum test, $p=3.6x10^{-6}$. **(B)** Mean SMYD3 mRNA expression levels in different cell types assessed from a publicly available single-cell RNA-seq database. Wilcoxon rank sum test with continuity correction, p<0.0001.

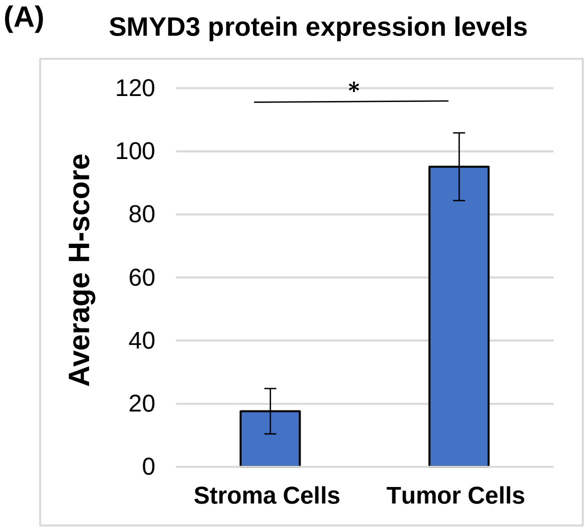


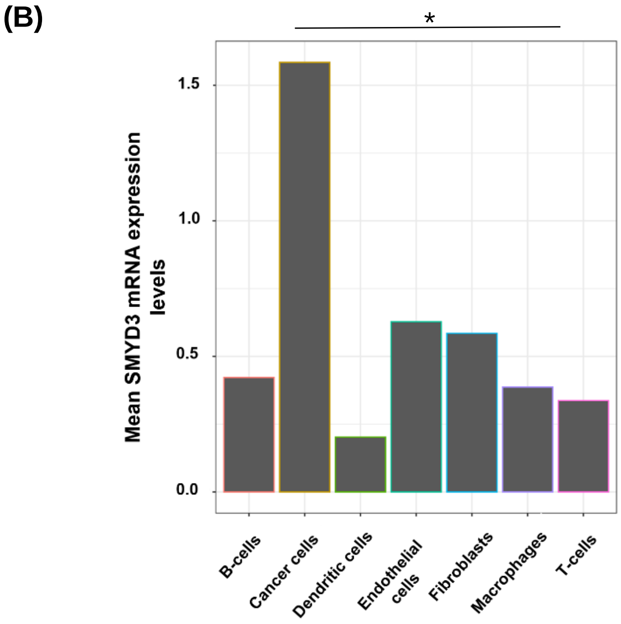

**Supplementary Figure 17.** Correlation between UHRF1 and CD8 protein levels in 64 HPV-negative HNSCC tumors. Pearson's correlation coefficient rho *R*=-0.011, p=093.

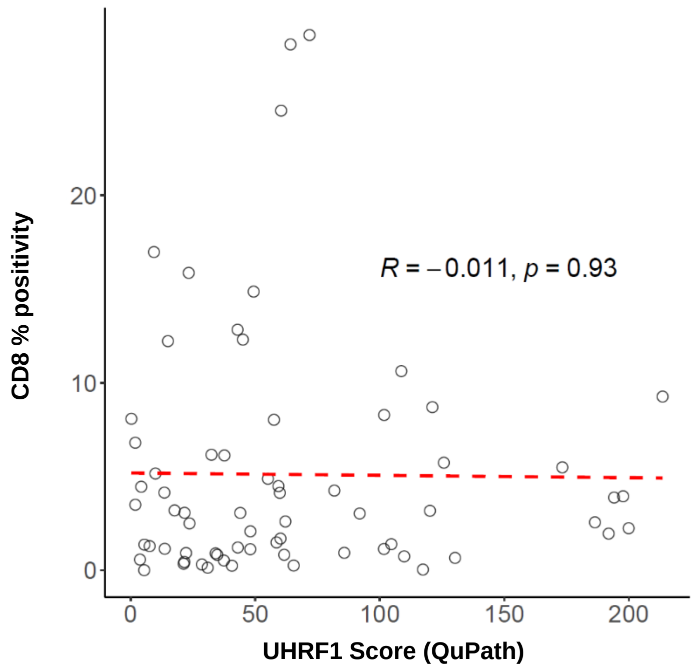

**Supplementary Figure 18.** SMYD3 (A), UHRF1 (B) QuPath Score and CD8 % positivity (C) and Kaplan Meier curves for overall survival.

**(A) SMYD3**

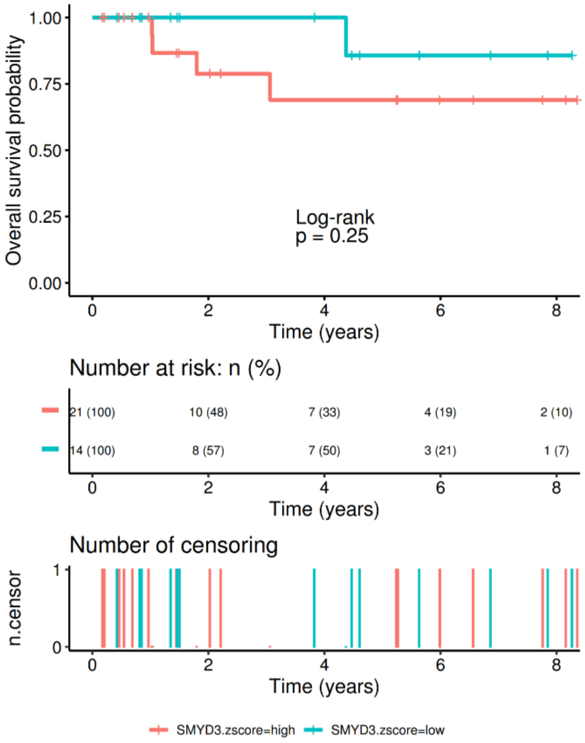


**(B) UHRF1**

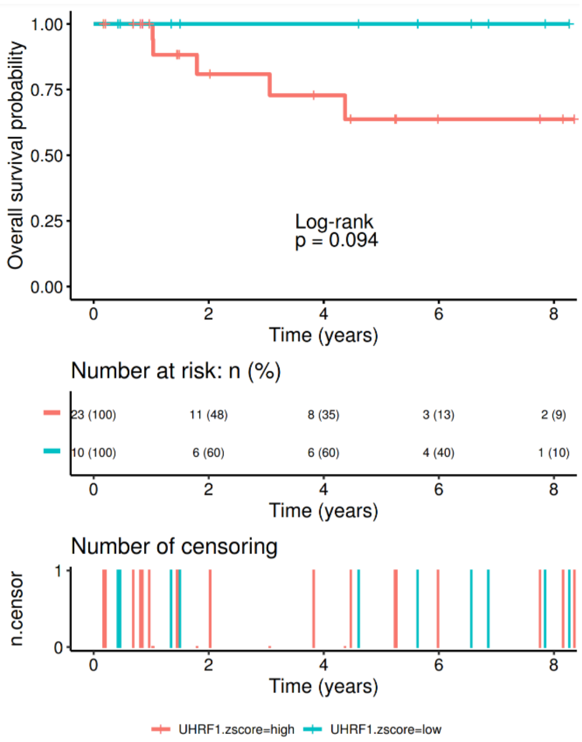


**(C) CD8**

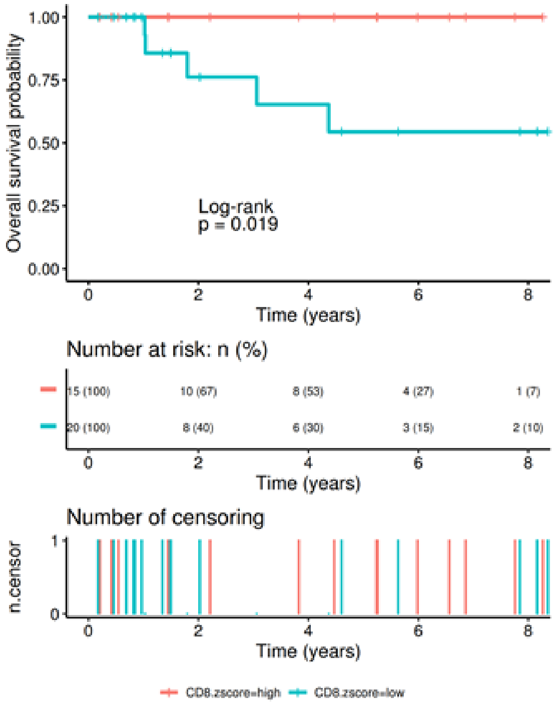

**Supplementary Figure 19.** mRNA expression levels of STING1 after SMYD3 depletion using siRNAs **(A)** or CRISPR (5-3 cell line) **(B)** and 24h of IFN-β exposure.

(A) HN-6 treated with siNC or siSMYD3 for 3 days

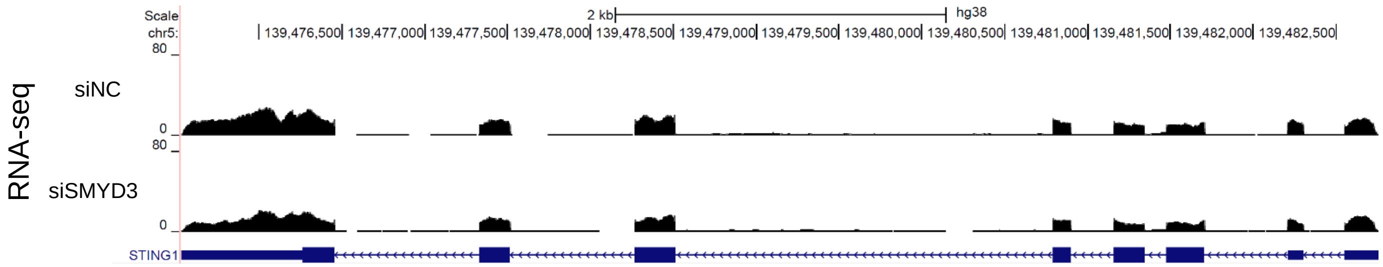


(B) SMYD3 KO 5-3 cell line

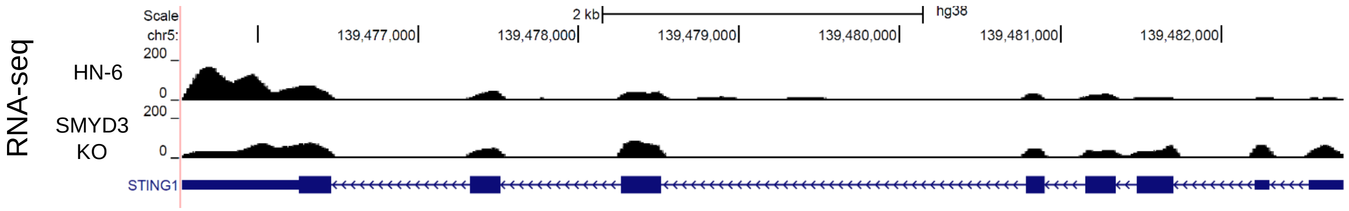


**Supplementary Figure 20.** Genomic alterations of SMYD3 in HPV-negative tumors samples of the TCGA dataset (Firehose Legacy).

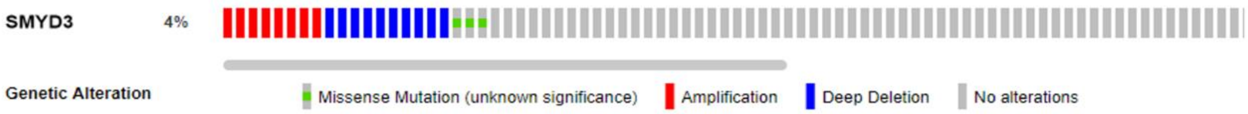


**Supplementary Figure 21.** Western blotting of SMYD3 in two CRISPR SMYD3 KO cell lines, 5-2 and 5-3. 10ug of nuclear extracts were loaded and blotted for SMYD3. H3 was used as a loading control.

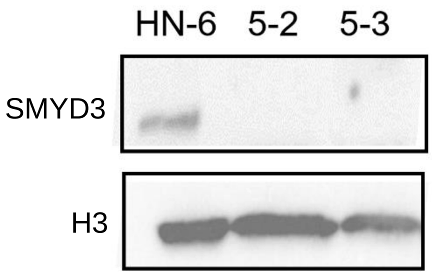
