## Supplementary Table 1 for "Epigenetic silencing by SMYD3 represses tumor intrinsic interferon response in HPV-negative squamous cell carcinoma of the head and neck"

**Supplementary Table 1. List of type I IFN response, APM and MHC class I genes**

**Type I IFN response genes**

CXCL9
CXCL10
TRIM21
MX1
MX2
RSAD2
DHX58
GBP2
CD274
OAS3
OASL
OAS2
STAT1
STAT3
IFNGR1
IL6
IL1A
IL1B
NCOA7

**APM and MHC class I genes**

TAP2
TAPBP
CANX
B2M
HLA-C
HLA-E
HLA-G
HLA-DMB
HLA-DPA1
HLA-DQB1
HLA-DRB1
HLA-DOB
HLA-DMA
HLA-DRA
