## Supplementary Table 4 for "Epigenetic silencing by SMYD3 represses tumor intrinsic interferon response in HPV-negative squamous cell carcinoma of the head and neck"

Supplementary Table 4. Immune-related genes with significantly decreased H4K20me3 peaks in promoters/TSS/gene body regions

| Number | Immune gene | Peak ID |
| --- | --- | --- |
| 1 | SAMD9 | P76842 |
| 2 | ELF1 | P18574 |
| 3 | IFIT3 | P10203 |
| 4 | RTP4 | P58964 |
| 5 | TENT5A | P71366 |
| 6 | TRIM25 | P32626 |
| 7 | SAMD9L | P76843 |
| 8 | DDX60 | P63993 |
| 9 | DDX60 | P63994 |
| 10 | HERC6 | P62297 |
| 11 | TDRD7 | P86672 |
| 12 | EPSTI1 | P18610 |
| 13 | IFI44L | P2619 |
| 14 | TENT5A | P71365 |
| 15 | TENT5A | P71358 |
| 16 | HLA-G | P69679 |
| 17 | LPAR6 | P18715 |
| 18 | TENT5A | P71360 |
| 19 | PROCR | P48537 |
| 20 | TENT5A | P71359 |
| 21 | TENT5A | P71356 |
| 22 | CREB1 | P45579 |
| 23 | PNPT1 | P42045 |
| 24 | HLA-DOB | P69767 |
| 25 | OAS1 | P17860 |
| 26 | HSP90AA1 | P24605 |
| 27 | GBP2 | P3688 |
| 28 | NFYC | P1119 |
| 29 | TENT5A | P71361 |
| 30 | HLA-B | P69730 |
| 31 | CMPK2 | P38569 |
| 32 | LPAR6 | P18714 |
| 33 | TMEM140 | P78149 |
| 34 | TDRD7 | P86676 |
| 35 | HSP90AA1 | P24606 |
| 36 | TENT5A | P71367 |
| 37 | CXCL11 | P61892 |
| 38 | TENT5A | P71363 |
